# A Rainbow Reporter Tracks Single Cells and Reveals Heterogeneous Cellular Dynamics among Pluripotent Stem Cells and their Differentiated Derivatives

**DOI:** 10.1101/2020.01.08.896290

**Authors:** Danny El-Nachef, Kevin Shi, Kevin M. Beussman, Refugio Martinez, Mary C. Regier, Guy W. Everett, Charles E. Murry, Kelly R. Stevens, Jessica E. Young, Nathan J. Sniadecki, Jennifer Davis

## Abstract

Recent single cell analyses have found molecular heterogeneities within populations of pluripotent stem cells (PSCs). A tool that tracks single cell lineages and their phenotypes longitudinally would reveal whether heterogeneity extends beyond molecular identity. Hence, we generated a stable Cre-inducible rainbow reporter human PSC line that provides up to 18 unique membrane-targeted fluorescent barcodes. These barcodes enable repeated assessments of single cells as they clonally expand, change morphology, and migrate. Owing to the cellular resolution of this reporter, we identified subsets of PSCs with enhanced clonal expansion, synchronized cell divisions, and persistent localization to colony edges. Reporter expression was stably maintained throughout directed differentiation into cardiac myocytes, cortical neurons, and hepatoblasts. Repeated examination of neural differentiation revealed self-assembled cortical tissues derive from clonally dominant progenitors. Collectively, these findings demonstrate the broad utility and easy implementation of this reporter line for tracking single cell behavior.

## Introduction

Single cell RNA sequencing studies have recently demonstrated that individual PSCs within a culture express a wide range of transcriptional profiles despite uniformly expressing the same cell-type specific markers, such as Oct4 and Tra1 (Kumar, Tan and Cahan, 2017; Nakanishi *et al*., 2019). It has been challenging to determine whether this translates to functional heterogeneity among PSCs. Most assessments of PSCs and their differentiated progeny are unable to track dynamic behaviors at the single cell level, presenting a technical limitation for the field. For instance, cell proliferation and morphology are typically quantified by staining for cell cycle or membrane markers in fixed specimens, which prevents repeated assessment of individual cells over time (Andäng *et al*., 2008; Kocsisova *et al*., 2018). Additionally, markers used for proliferation studies are not definitive indices of newly generated cells (van Berlo and Molkentin, 2014). For example, PSC-derived cardiac myocytes and hepatocytes can undergo DNA synthesis and nuclear division without completing cell division (Hannan *et al*., 2013; Lundy *et al*., 2013). Live-cell markers, such as GFP fusion proteins with cell cycle or membrane markers, have been used to overcome this limitation (Gantz *et al*., 2012; Pauklin and Vallier, 2013), but this approach is still hampered by the inability to resolve the parental identity of cell progeny or distinguish differences in clonal expansion since the cells are uniformly labeled with the same color. Likewise, studies of PSC and PSC colony migration are challenging due to difficulty in following them over long periods of time in the absence of live-cell barcodes (Kusuma and Gerecht, 2013; Megyola *et al*., 2013; Zangle *et al*., 2013). Since PSCs have emerged as important tools to regenerate tissue, model human development and disease, and for drug screening (Inoue *et al*., 2014; Rowe and Daley, 2019), realizing the dynamic behaviors of PSCs on a per cell basis could improve our understanding of human biology and medicine.

Recently, rainbow mouse models have enabled tracking of single cell lineages (Livet *et al*., 2007; Snippert *et al*., 2010). In these models, Cre recombinase (Cre) treatment induces the expression of a combination of fluorescent proteins, which in turn barcodes cell lineages with a broad palette of hues. Because the DNA recombination is permanent, the color barcode of a cell is inherited by daughter cells during cell division, enabling determination of the parental origin of cellular progeny. The first of these models utilized neuron-specific promoters to express the rainbow labels, i.e. Brainbow (Livet *et al*., 2007; Cai *et al*., 2013). A confetti mouse model was also engineered to extend studies to non-neuronal cell types by exchanging the Thy1 neuron-specific promoter to a ubiquitous CAG promoter upstream of a modified Brainbow cassette containing three unique fluorescent proteins in tandem (Snippert *et al*., 2010). A limitation of the Confetti model is the color reporters are differentially expressed by Cre-mediated inversion rather than deletion that can cause a single cell to change colors multiple times rather than maintain a permanent label. Both models are also limited in their applicability to cell morphology studies because some of the fluorescent reporters lack membrane targeting. A more recent version, Brainbow 3.2, permanently barcodes neurons with fluorescent proteins that are all targeted to the plasma membrane, thereby facilitating lineage tracing, delineation of individual cells, and morphological analysis (Cai *et al*., 2013). While this fluorescent lineage technology has been utilized extensively *in vivo*, to our knowledge it has not been applied to human stem biology. Indeed, rainbow stem cell lines would permit repeated assessment of the same cell and its progeny over long periods of time *in vitro*, enabling assessment of cell dynamics, such as rates of clonal expansion, cell cycle kinetics, and synchronization of cell divisions which cannot be assessed with the current *in vivo* rainbow technology. In addition, membrane-targeted fluorescent signals could be leveraged to measure temporal changes in cell morphology.

Here we demonstrated that engineering a knock-in PSC line with the Brainbow 3.2 cassette under the control of a CAG promoter enabled repeated live-imaging of fluorescently barcoded PSCs and robust quantification of cell dynamics during PSCs self-renewal and differentiation into multiple cell types. Moreover, directed differentiation of the rainbow PCSs into cortical neurons uncovered that three dimensional (3D) cortical structures are derived from clonal expansion of dominant progenitors, a finding made possible by employing the rainbow reporter stem cell line. In total this work reports the generation of the first, to our knowledge, rainbow human PSC lines and demonstrates the broad utility of this tool to track single cell behavior and clonal expansion of lineages in real-time and throughout directed differentiation.

## Results

### Engineering rainbow knock-in stem cell lines

To generate a universal rainbow cell reporter in the WTC11 human induced-PSC (hiPSC) line we targeted the AAVS1 safe-harbor locus and knocked-in a cassette containing a constitutively active CAG promoter upstream of three distinct fluorescent proteins (eGFP, mOrange2, and mKate2, hereafter referred to as GFP, RFP, and FRFP) with each possessing a stop-codon and flanked with incompatible lox sequences for Cre-mediated recombination (**Figure 1A**). The construct is designed such that a non-fluorescent nuclear-localized GFP-mutant (nFP) is constitutively expressed until Cre treatment, which initiates permanent recombination of the construct. These recombination events cause one of three fluorescent proteins to be expressed on the cell membrane that is passed on to its progeny. Targeted knock-in was confirmed and lines with multiple copies of the cassette were generated (**Figures S1A and S1B**), allowing for the expression of unique color combinations in individual cells, which act as a visible fluorescent barcode (**Figure 1B**). We selected a knock-in line with 4 copies of the rainbow cassette for further study as it permits the expression of 18 unique hues (**Figure 1B**).

**Figure 1.**
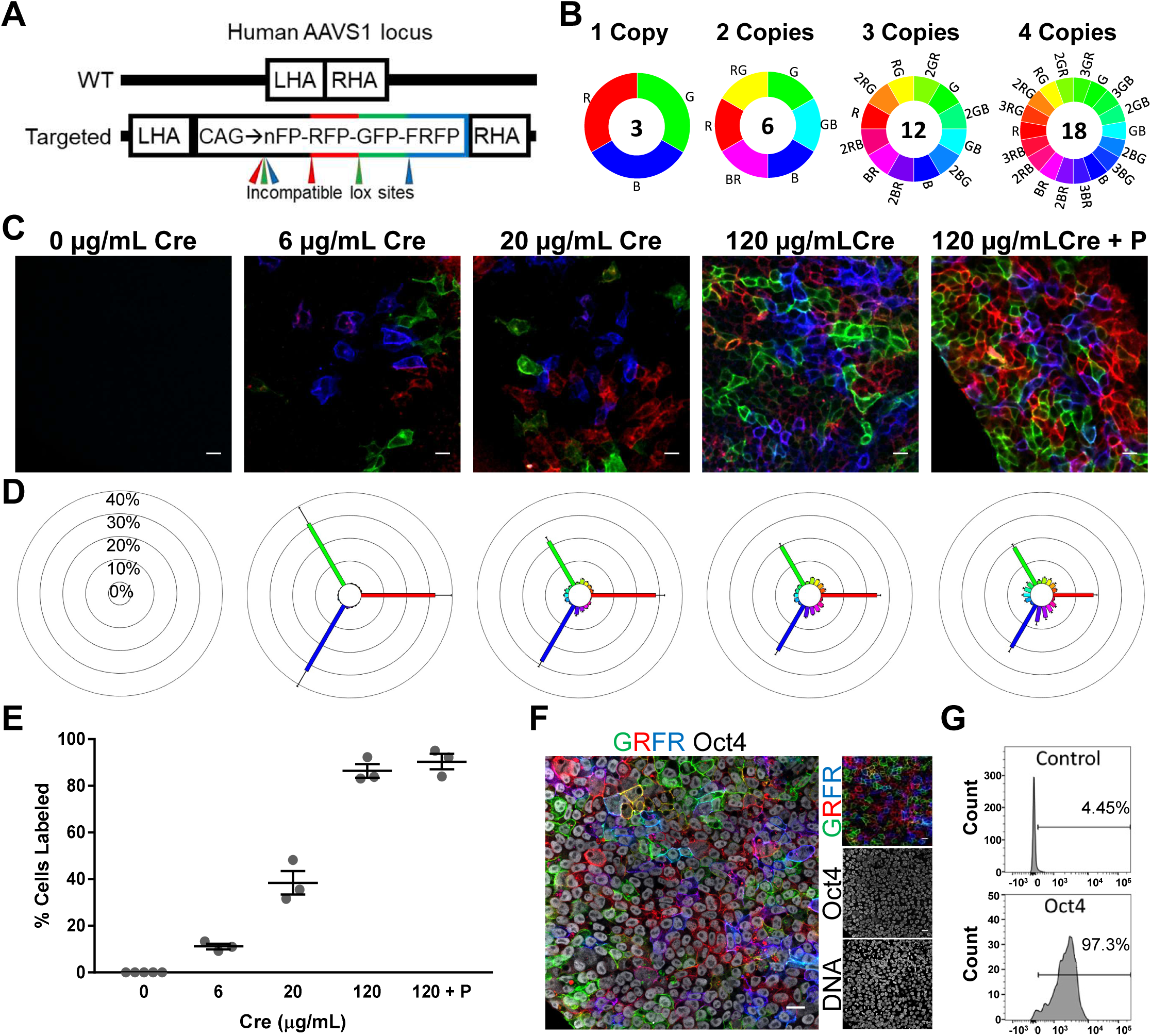
Authentication of rainbow hiPSC model. (A) Schematic of the human AAVS1 safe-harbor locus depicting the wildtype allele with left homology arm (LHA) and right homology arm (RHA) (top) and targeted, knocked-in rainbow cassette containing GFP, RFP, and FRFP flanked by incompatible lox sites for Cre-mediated recombination (bottom). (B) The number of unique hues possible for a given number of copies of the rainbow cassette. Recombination outcomes that result in the same hue, i.e. recombination of one cassette that expresses RFP versus two cassettes that both express RFP, were considered one unique hue. (C) Representative confocal images of rainbow hiPSCs that were treated with various doses of Cre and imaged using white light laser and spectral detector settings that prevented spectral bleedthrough of the fluorescent proteins. Cells were imaged 72 hours after Cre treatment (0-120 µg/mL), or after more than 8 passages (120 µg/mL + P). Bar=20 µm. (D) Spectral analysis of the percentage distribution of hues from the various Cre doses. (E) Proportion of cells expressing at least one fluorescent protein with varying Cre doses. (F) Native rainbow fluorescence and immunofluorescence imaging of Oct4. Bar=20 µm. (G) Representative trace of flow cytometry analysis of Oct4+ rainbow hiPSCs, controls were treated equally but without addition of primary antibody to Oct4.

Rainbow hiPSCs were treated with different doses of recombinant cell-permeant Cre to vary fluorescent protein expression. The proportion of labeled cells and diversity of color barcodes were assessed 72 hours or several weeks (passaged) after Cre treatment (**Figures 1C and S1C**). Using confocal microscopy with white light laser and spectral detector settings that prevented bleed-through of fluorescent signals demonstrated that the proportion of labeled cells and color diversity could be titrated by varying Cre dose. A low Cre dose of 6 µg/mL resulted in recombination of only a single rainbow cassette as demonstrated by the expression of primary colors (red, green, and blue) at equal percentages of labeled cells (**Figure 1D**). High Cre doses induced the expression of additional hues, which develop via the expression of more than one fluorescent protein yielding cells with yellow, purple, or turquoise, and a reduced percentage of primary colors (**Figure 1D**). Reporter expression was stable in cells that had been passaged eight or more times after initial Cre treatment with the percent of labeled cells and the distribution of hues remaining consistent after many passages (**Figures 1D and 1E**), indicating there was no selection against labeled cells and that no particular color had a proliferative advantage on a cell population level.

Rainbow hiPSCs retained their stemness, expressing stem cell markers Oct4 (**Figures 1F and 1G**), and Tra1 (**Figure S1D**), and did not have chromosomal abnormalities in their karyotype (**Figure S1E**). Immunostaining with antibodies that can distinguish each fluorescent protein as well as nFP showed individual cells that expressed all the fluorescent proteins and lacked nuclear-localized nFP expression, indicating all four rainbow cassettes could be recombined when treated with high Cre dose (**Figure S2**). The immunostaining and choice of secondary antibodies also allowed us to visualize the spectral diversity of the cells using standard filter cubes (**Figure S2**). A rainbow reporter in the RUES2 human embryonic PSC line was validated using the same approach (**Figure S3**).

### Tracking single cell proliferation kinetics reveals hiPSCs synchronously divide

To confirm our rainbow line was a suitable reporter of cell proliferation, immunostaining of phosphorylated histone H3 was performed to quantify the percent of mitotic cells in isogenic wildtype WTC11 and rainbow knock-in hiPSCs after high-dose Cre treatment. Analysis at 24, 48, and 72 hours after plating indicated there were no differences in proliferation between Cre-treated multicolored rainbow hiPSCs, rainbow hiPSCs with no Cre treatment, or non-transgenic control hiPSCs at any timepoint (**Figures S4A and S4B**). In addition, analysis of TUNEL positivity at 72 hours post seeding showed no differences in apoptosis between these groups (**Figure S4C**).

Rainbow hiPSCs were live imaged and tracked over 24 hours (**Figure 2A and Videos 1-3**) to examine hiPSC proliferation in real time and quantify cell cycle kinetics. Because individual hiPSCs were distinguishable by the rainbow barcodes and images were acquired with fine temporal resolution that captured cytokinesis events, we were able to follow cell division as a parent cell divided into two daughter cells (D1 and D2) and continue tracking subsequent cell divisions of daughters into granddaughter cells (*e*.*g*. D1.1 and D1.2, **Figure 2B**). Tracking cell cycle duration of daughter cells, as well as tracking cells that originated from the same progenitor at the time of plating, showed that cell cycle duration in these groups had no differences compared to all cells examined (**Figure 2C**). To our knowledge, this analysis is the first to report hiPSC proliferation kinetics in which the average cell cycle duration was 13.6 ± 0.14 hours, which is consistent with other reports in human and mammalian embryonic stem cells that estimated 11-16 hour cell cycle lengths (Orford and Scadden, 2008).

**Figure 2.**
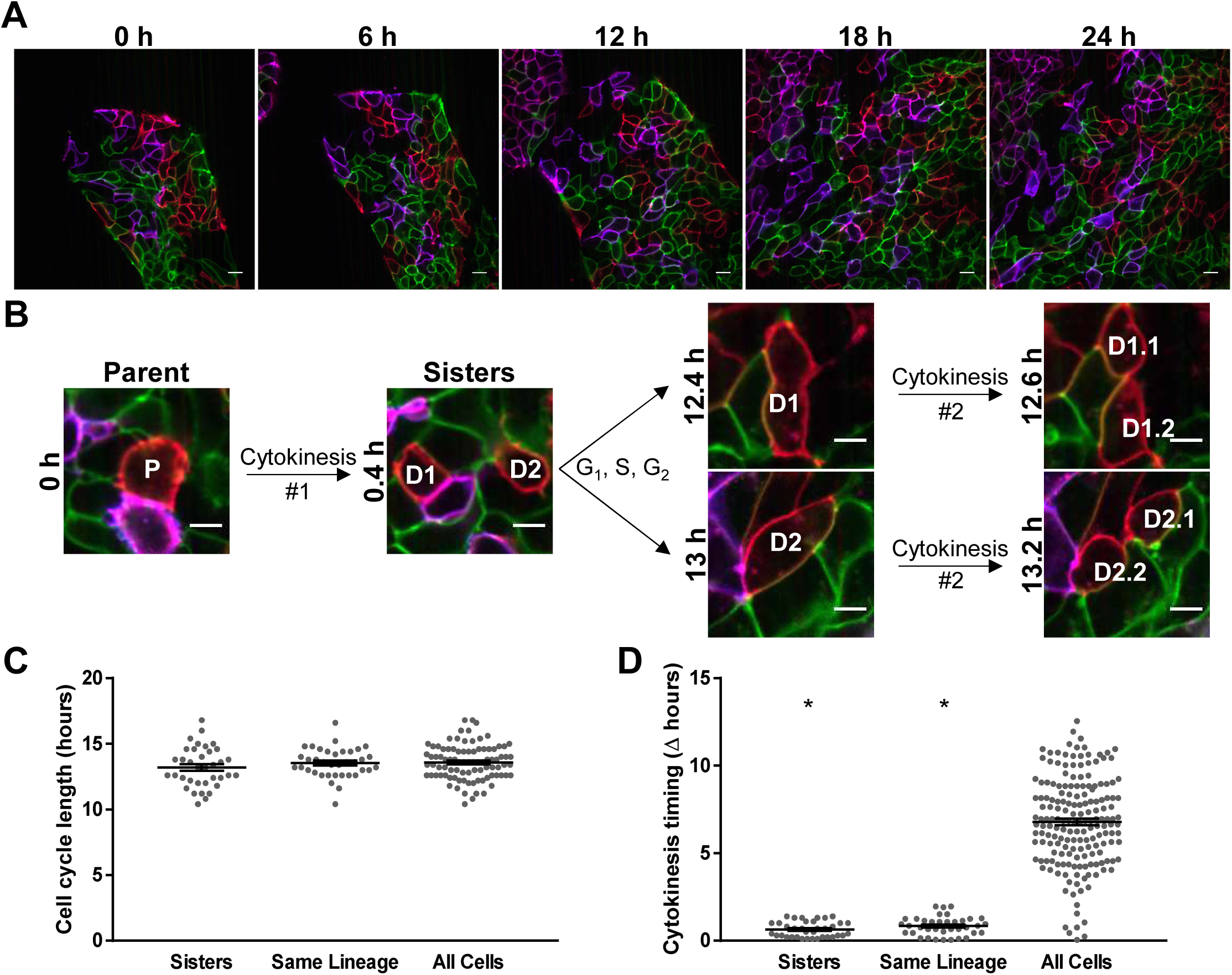
Real time, longitudinal assessment of proliferation kinetics in live hiPSCs reveals synchronized cell divisions. (A) Representative images of 24-hour time-lapses (121×12 minute frames spanning the period of 3-4 days after plating, related to Videos 1-3). One frame every six hours during a time-lapse is shown, bar=20 µm. (B) Representative example of tracking single parental cell (P) just before it divides (t=0 hr), its daughters (D1 and D2 sisters), and granddaughters (D1.1, D1.2, D2.1, and D2.2), bar=10 µm. (C) Quantification of cell cycle duration of tracked sister cells, cells from the same lineage (common ancestor cell at time of plating), and all cells measured. (D) Difference from mean cell division timepoint among sister cells, cells from the same lineage, and all cells. * indicates p < 0.05 versus all cells.

A common feature of early metazoan embryogenesis, including mice and humans, is synchronized cell division in PSCs *in vivo* (O’Farrell, Stumpff and Tin Su, 2004; Wong *et al*., 2010; Meseguer *et al*., 2011; Desai *et al*., 2014; Deneke *et al*., 2016). In contrast, *in vitro* studies of mammalian PSCs have suggested PSC divisions are not synchronized, unless one manipulates the cell cycle machinery with chemical inhibitors or serum deprivation (Zhang *et al*., 2005; Yiangou *et al*., 2019). However, tracking the timing of cell divisions in progeny that arose from a single PSC demonstrated that PSCs arising from the same lineage underwent synchronous cell divisions (**Figure 2D**). We observed that sister cells divided within 0.64 ± 0.10 hours of each other and cells of the same lineage divided within 0.78 ± 0.16 hours. In contrast, any given cell divided within 6.79 ± 0.19 hours of the average division of the population, consistent with the apparent asynchronous behavior observed *in vitro* (Zhang *et al*., 2005). Lastly, of all the cells examined, we observed that 2.7% ± 0.9% underwent apoptosis (**Figure S4D**), which is consistent with results using TUNEL quantification (**Figure S4C**). These results demonstrate cell cycle kinetics and synchronization of cell divisions can be quantified with the rainbow reporter.

### Long term, longitudinal assessment of live hiPSCs reveals heterogeneous clonal expansion

To extend the longitudinal assessments to periods lasting more than a week, hiPSCs were imaged 2 hours after sparsely seeding (day 0) and then repeated imaging occurred once every 24 hours for 7 days (**Figures 3A and 3B**). To prevent distinct clones of the same hue from becoming near each other due to migration and/or proliferation, rainbow hiPSCs treated with 120 μg/mL Cre to achieve a high color diversity were mixed with rainbow hiPSCs that were not treated with Cre at a 1:10 ratio. The number of cells per clone was quantified at each time point (**Figure 3C**). On average the number of cells per clone increased from 1.2 cells at plating (day 0) to 324.4 cells after a week of culture (p < 0.0001). The greatest rate of clonal expansion occurred at 3 to 4 days after plating, which then slowed as the cultures became confluent (p < 0.05 versus rate of expansion day 6 to 7, **Figure 3C**). Vast differences between the expansion rate of individual clones was observed. For instance, two individual cells plated on day 0 expanded to colonies containing 53 and 1054 cells by day 7, which indicates a clonal expansion rate of 1.87-fold increase per day versus 3.13-fold per day (**Figure 3D**). These results suggest that mechanisms intrinsic to the cells or cues arising from their microenvironment, such as cell crowding, have effects on clonal expansion. On a cell population level however, no particular hue or color combination of cells had a proliferative advantage, as demonstrated by the consistency of color distribution after more than 8 passages (**Figure 1D**). This analysis highlights how heterogeneous clonal expansion can be uncovered using live-cell rainbow barcodes.

**Figure 3.**
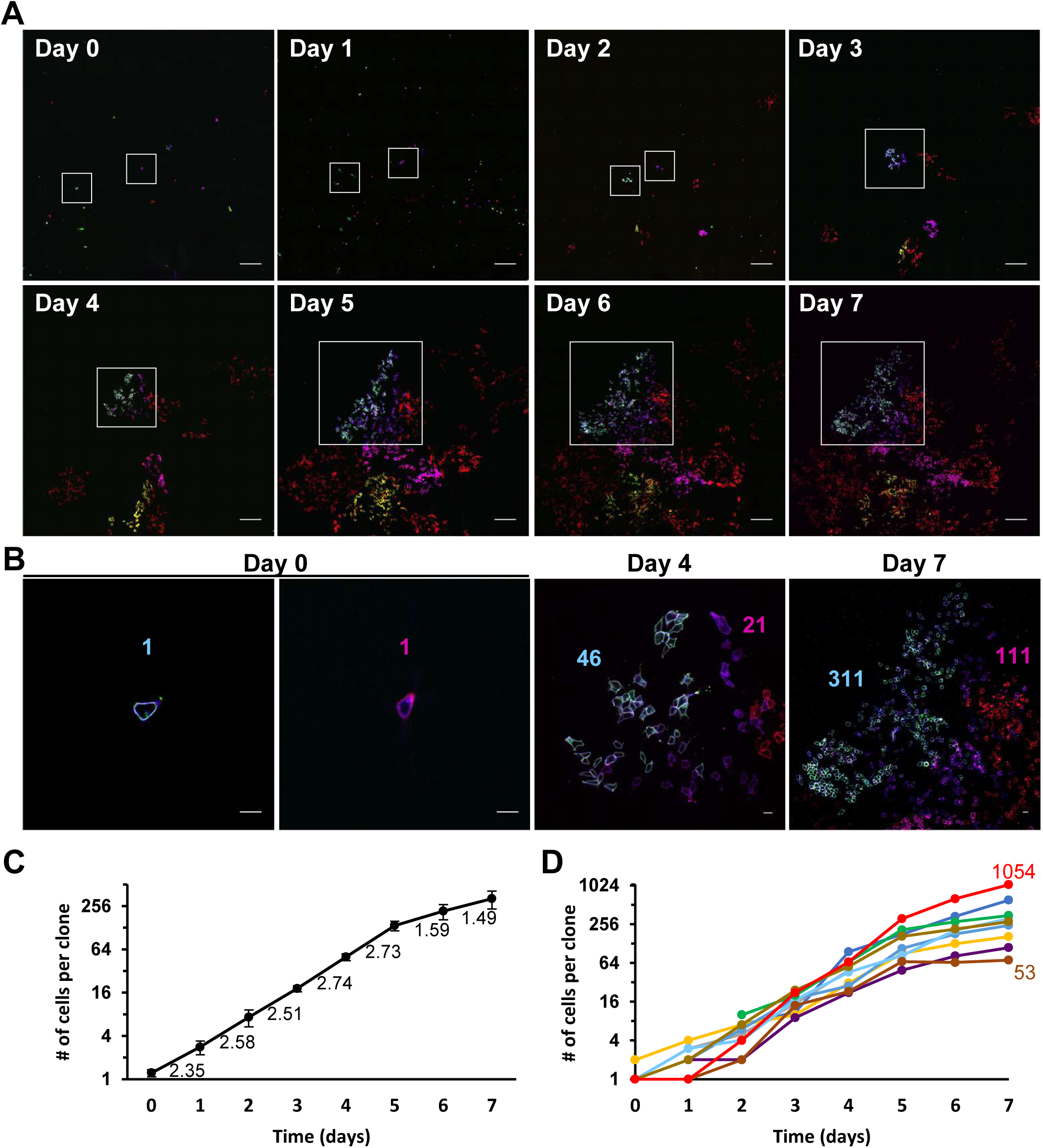
Long term, longitudinal assessment of live hiPSCs reveals heterogeneous clonal expansion. (A) Labeled cells were mixed 1:10 with nonlabelled transgenic hiPSCs, seeded sparsely, and repeat imaged once every 24 hours for 8 days on a live-imaging spinning disk confocal microscope. Day 0 = 2 hours post plating and all following timepoints are in 24-hour intervals, bar=200 µm. (B) Single cell resolution images of two clones with cell number denoted in the same color as respective clone at day 0, day 4, and day 7 post seeding, bar=20 µm. (C) Quantification of average clonal expansion over the course of culture, fold increase in cell number per clone indicated below line. (D) Cell number per clone plotted individually for each clone, highlighting divergent clonal expansion.

### hiPSCs undergo dynamic morphogenesis

To understand how cell morphology changes during a cell cycle, time-lapse datasets were analyzed (**Figure 2 and Videos 1-3**) and the membrane-localized fluorescent signal was traced throughout the cell cycle (**Video 4**). Newborn cells hypertrophied through the cell cycle, being largest just before cell division (**Figure 4A**), consistent with previous reports in mammalian cells (Lloyd *et al*., 2000; Girshovitz and Shaked, 2012). The boundaries of cells in the previous dataset (**Figure 3**) were traced to determine how their area changed 1 week in culture. Cell area decreased with confluency (**Figure 4B**) as indicated by 390.8 µm^2^ at day 0 versus 134 µm^2^ at day 7 (p < 0.0001). However, examination of confocal Z-stacks (**Figure 4C and Video 5**) showed that as cell become smaller in their lateral dimensions they grow taller in height (9.2 µm at day 0 versus 38.1 µm at day 7, p < 0.0001) (**Figure 4D**), demonstrating hiPSCs do not get smaller when confluent but rather change shape.

**Figure 4.**
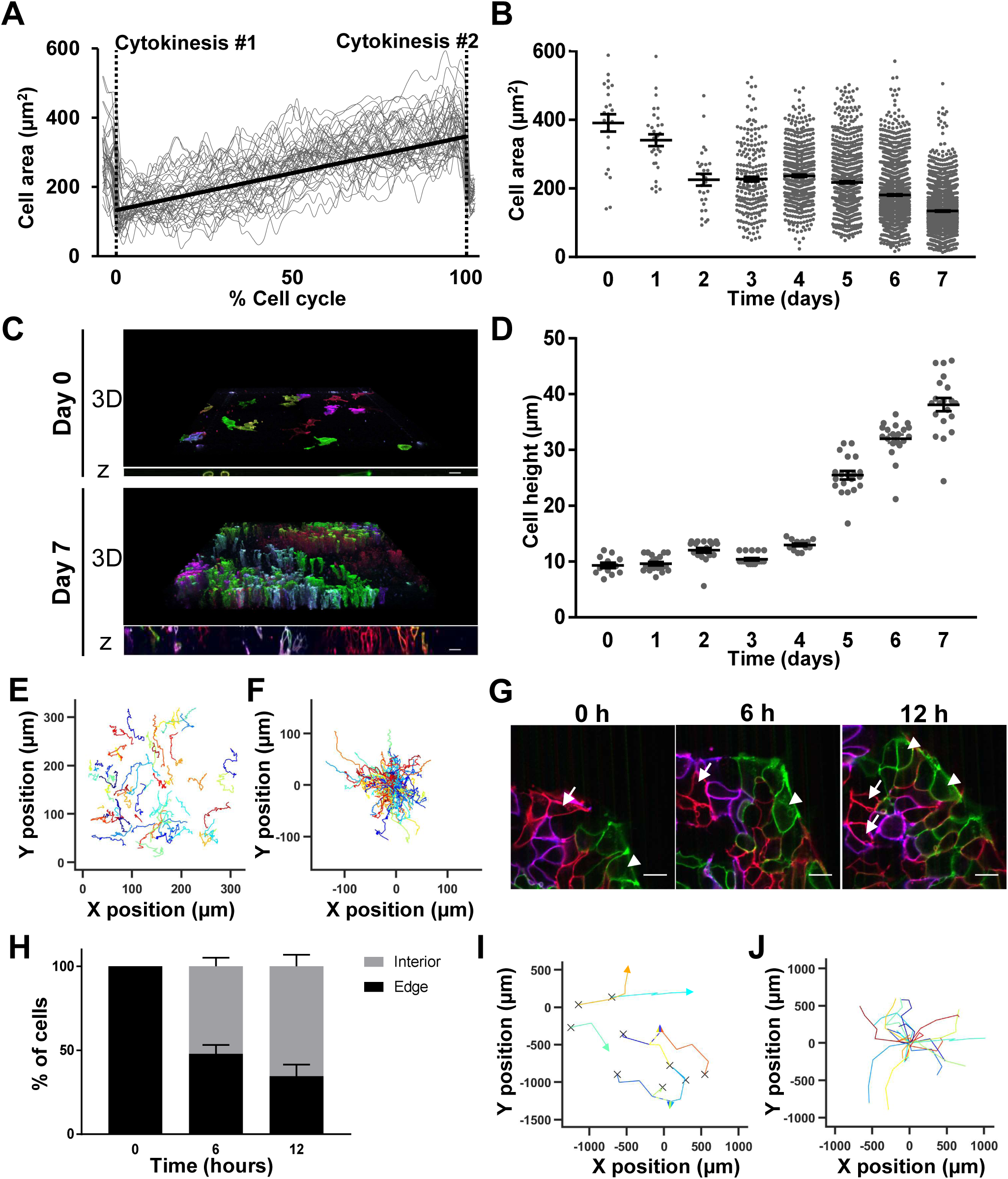
Longitudinal assessment of hiPSC morphogenesis reveals hypertrophy in cell cycle growth phase, confluency-induced shape changes, and colony migration. (A) Cell area measurements spanning two cell divisions with cell area measured every 12 minutes. The duration of each cells’ cell cycle length was normalized to % cell cycle. Gray lines indicate individual cells, black line indicates line of best fit, and dashed vertical lines indicate cytokinesis time points at 0% and 100% of the cell cycle. (B) Cell area measurements over 8 days in culture. (C) Examples of 3D reconstructions at Day 0 and Day 7 post plating (601µm x 601µm, 0.4µm steps), bar=20 µm. (D) Quantification of cell height over 8 days. (E and F) Migration analysis tracking absolute (E) or relative (F) positions of individual hiPSCs. (G) Representative image showing an example of one parental cell on the colony perimeter at 0 hours that gave rise to two internally localized cells at 12 hours (white arrows) and another that gave rise to two cells with maintained peripheral localization (white arrowheads), bar=20 µm. (H) Analysis of edge PSCs showing proportion remaining on the edge or translocating interiorly at each time point. (I and J) Migration analysis tracking absolute (I) or relative (J) hiPSC colony centroid position from Day 0 to Day 4. (I) “X” denotes colony start position and arrowheads indicate colony end position. Colony fusion events are indicated by dashed lines and split-color arrowheads.

The rainbow reporter also enabled delineation of individual hiPSCs within densely packed colonies, which revealed a highly migratory behavior. Their migration was assessed by tracking the position of the cells over time (**Figure 4E and Videos 1-3**) and quantifying the overall distance traveled and direction of migration (**Figure 4F**). The migration behavior was different between the tracked cells, with velocities ranging from 7.3 to 20.3 µm/hour. Some hiPSCs located on the colony’s edge could translocate interiorly while others remained at the edge during colony expansion (**Figure 4G**). Tracking spatial locations of a cell within a colony can yield new information regarding geometric cues, as recent work identified a subpopulation of PSCs on colony edges that share hallmark properties with primitive endoderm, with distinct cell states compared to those in the interior (Yoney *et al*., 2018; Nakanishi *et al*., 2019). To understand the kinetics of PSC translocation from the perimeter to the interior, PSCs at the edge of colonies were tracked over a period of 12 hours and analyzed for position changes from the edge to the interior (**Figure 4H**). Over this observation period, 84% of the cells divided and none underwent apoptosis, so all the cells starting at the colony edge and their progeny were able to be tracked. By 6 hours, 47.9% ± 4.3% of the cells that started at the edge were still localized there (p < 0.001 versus 0 hours) and by 12 hours, it dropped to 34.6% ± 5.6% of the cells (p < 0.001 versus 0 hours) (**Figure 4H**). Of the cells that migrated to the interior after 6 hours, only 4.2% ± 3.4% migrated back to the colony edge in the subsequent 6 hours, suggesting that hiPSCs dynamically migrate within PSC colonies and translocate from the periphery to the interior compartment. The rainbow stem cell colony-centers migrated 211.7 µm ± 12.9 µm per day (**Figures 4I and 4J**). Colony fusion events over long periods of culture were clearly distinguishable (**Figures 4I and 4J**), including the formation of colonies that arose from multiple, distinct fusion events. Together, these results demonstrate the membrane-specific labeling can be exploited to assess the morphogenesis of rainbow barcoded cells and colonies.

### Rainbow hiPSCs are pluripotent and maintain reporter expression throughout differentiation to all germ layers

Rainbow hiPSCs were differentiated to cortical neurons (ectoderm), cardiac myocytes (mesoderm), and hepatoblasts (endoderm) (**Figure S5A**) as defined by the expression of key proteomic markers (Gantz *et al*., 2012; Mariani *et al*., 2012; Mallanna and Duncan, 2013) (**Figures 5A-5C**). These data demonstrate that the rainbow reporter is maintained during directed differentiation to all three germ layers and innocuous with respect to their pluripotency (**Figures 5A-5C**). Because neurogenesis encompasses clonal expansion, cell migration, and dramatic morphological changes (Lancaster *et al*., 2013), we examined whether these behaviors could be tracked with the rainbow reporter in differentiated neurons. Here images were acquired daily during the first weeks of neurogenesis as hiPSCs committed to the neural lineage (**Figures S5B-S5F**). Tracking individual cells and repeat imaging of their progeny provided evidence of clonal expansion during the formation of Sox2+ neuronal progenitor cells (NPCs) (**Figures S5B and S5C**). Subsequent live imaging demonstrated clonal expansion and dispersal of NPCs as they elongated and radially aligned to form neural rosettes, which were similar in structure compared to the isogenic wildtype control (**Figures S5D-S5F**). Similarly, cells were repeat imaged during the differentiation of rainbow hiPSCs to cardiac myocytes, demonstrating clonal expansion of cardiac progenitors, maturation as indicated by hypertrophy, and high levels of cardiac troponin T expression (**Figures 5D and S5G**). Functional assessment of rainbow-labeled hiPSC-derived cardiac myocytes demonstrated that contractions could be visualized by imaging deformations of the rainbow-colored membranes at a high frame rate (**Video 6**). Additionally, the rainbow reporter could be activated with temporal control based on the timing of Cre delivery, as shown by labeling cardiac myocytes after unlabeled rainbow hiPSCs underwent differentiation (**Figure S5H**). These findings confirm the rainbow reporter can broadly assess clonal expansion and morphogenesis longitudinally in differentiated cells and hiPSCs.

**Figure 5.**
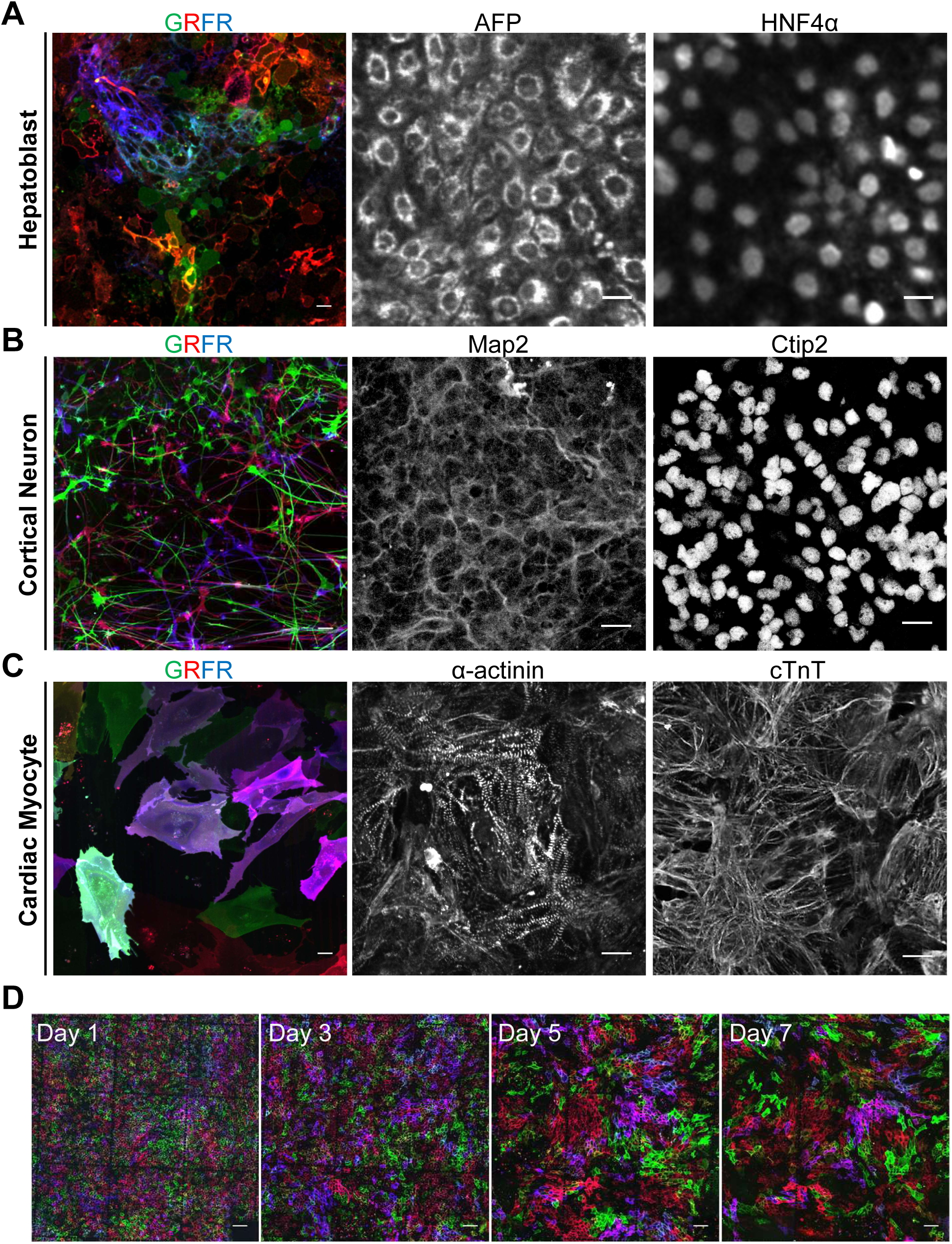
Rainbow hiPSCs have capacity to differentiate to all germ layers and maintain reporter expression. All differentiations started with high dose Cre-treated rainbow hiPSCs replated at Day 0. Rainbow reporter signal and respective cell-type specific marker expression in (A) hepatoblasts, (B) cortical neurons, and (C) cardiac myocytes, bars=20 µm. (D) Repeat imaging of rainbow labeled hiPSCs to CMs starting from the first day of differentiation through one week (Days 1-7), bars=100 µm.

### Self-assembled 3D cortical structures are derived from clonally dominant progenitors

Since the rainbow reporter was stable and could be visualized over week-long periods of differentiation, time-lapse analysis studies were extended to assess the later phases of neurogenesis that encompass cortical neuron formation and terminal differentiation. After 24 days of NPC differentiation and expansion, NPC cultures were dissociated into single cells, replated, and repeat-imaged over the subsequent 30 days of cortical neuron differentiation. By day 54 a few clonally dominant NPCs had generated most of the cells in the cultures, as evidenced by large clusters harboring the same fluorescent barcode (**Figures 6A and 6B**). Assessment at 3 days after replating (day 27) demonstrated homogeneous dissociation and mixing of cells that expressed a broad range of color barcodes (**Figures 6A and 6B**), confirming that clonal dominance was not driven by initially seeding large clusters of same-colored NPCs. Time-lapse images showed expansion of a subset of clones, further confirming this finding (**Figures 6A and 6B**). The day 54 culture was primarily neurons, though we noted DCX^-^ Map2^-^ cells at the center of massive clonal aggregates that had formed (**Figures 6A and 6B**). Confocal imaging revealed these clonal aggregates were three-dimensional and had multiple cell layers, reaching greater than 100 µm in height (**Figure 6C**).

**Figure 6.**
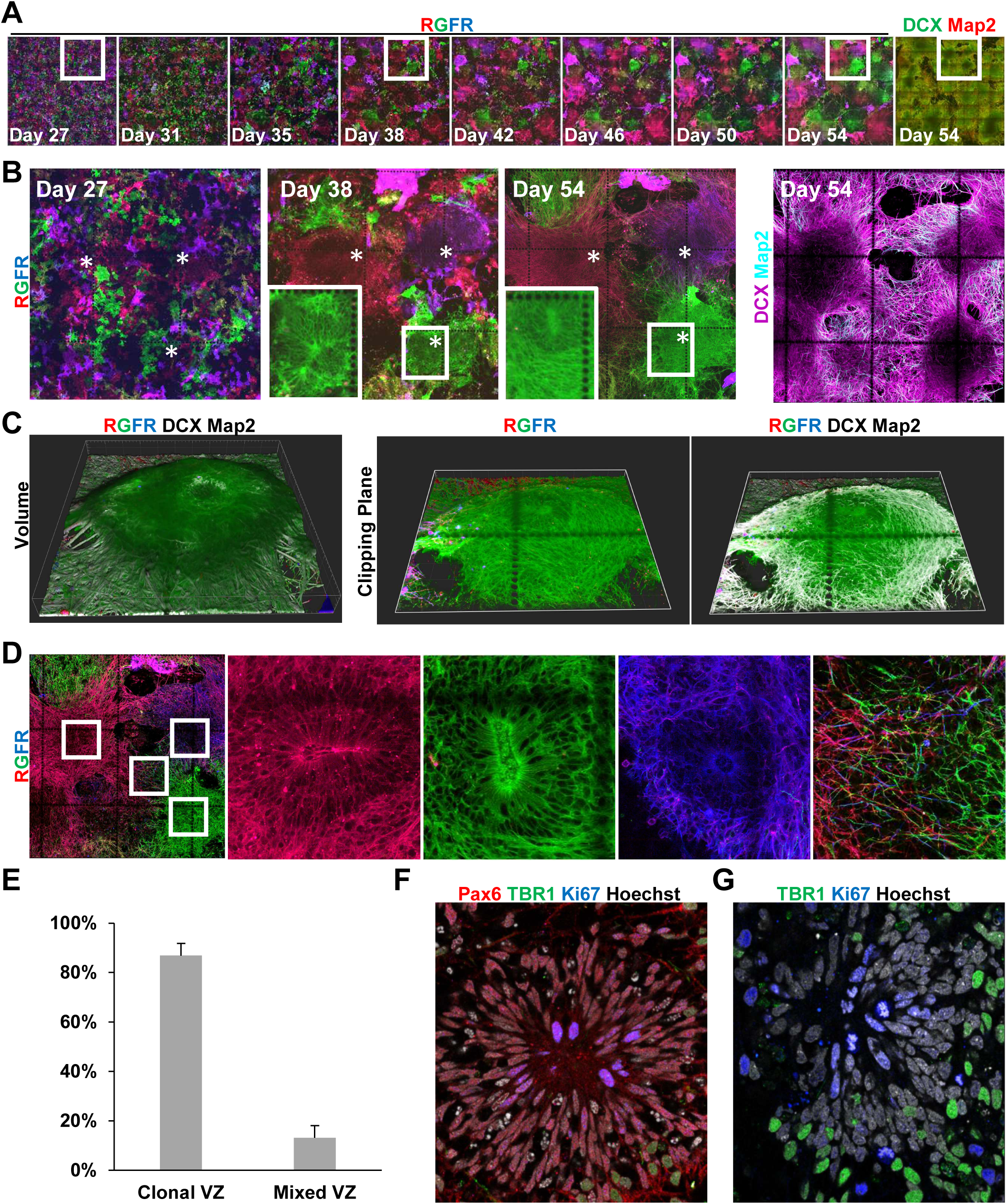
Self-assembled 3D cortical structures are derived from clonally dominant progenitors. (A) Repeat imaging of the rainbow label in NPCs differentiating to cortical neurons, and immunostaining confirmation of neuronal differentiation at the Day 54 endpoint. (B) Higher magnification of selected timepoints from panel A. Asterisks indicate the starting position of three clones that were dominant. Insets are higher magnifications of different z-planes of the boxed regions, localized 16 µm above the culture substrate to visualize VZ structure. (C) Day 54 3D reconstructions of the entire z-stack volume and clipping planes through the volume. (D) Examples of clonal VZs with insets showing higher magnifications of different z-planes of the boxed regions highlighting VZ structure and distal neurite outgrowth. (E) Quantification of the proportion of VZs derived from a single NPC (Clonal VZ) or VZs derived from multiple NPCs. (F and G) examination of cell cycle activity in NPCs and terminally differentiated cortical neurons. (A-G) Each coordinate grid represents a 500 × 500 µm square and each dot has a 10 µm diameter.

To better understand the formation and identity of cells in these 3D structures, confocal z-stacks over the live imaging time course were examined. The membrane-targeted label revealed that by Day 38, NPCs in the monolayer culture had self-organized into radially aligned cells forming a ventricular zone (VZ) in these aggregates (**Figure 6B inset**). Imaging at subsequent timepoints showed neurites extending out from the VZs (**Figure 6B and Video 7**). These features closely resemble neurogenesis in the developing neural tube and telencephalon *in vivo* (Graham *et al*., 2003; Moreno and González, 2017), as well as cortical organoids generated by PSC differentiations in 3D droplets (Lancaster *et al*., 2013; Paşca *et al*., 2019), suggesting monolayer differentiations also result in the formation of complex cortical structures. Immunostaining at Day 54 confirmed the NPC identity of the radially aligned cells persisted in the VZs, staining positive for Sox2 and Pax6 (**Figures S6A and S6B**). Analysis of the rainbow reporter and neurite markers at day 54 demonstrated that NPCs in VZs generated neurons expressing the same color barcode, with newborn DCX^+^ neurons residing proximal to the VZ and MAP2^+^ terminally differentiated neurons localizing more distal to the VZ (**Figures 6B, 6C and S6A**), consistent with neurogenesis *in vivo* (Moreno and González, 2017; Johnson *et al*., 2018). Co-staining of Pax6 and TBR1 (**Figure S6C**) confirmed the self-organized spatial patterning of NPCs and postmitotic cortical projection neurons *in vitro* were consistent with mammalian corticogenesis *in vivo* (Englund *et al*., 2005; Bedogni *et al*., 2010).

Surprisingly, of the 183 VZs examined at day 54, 89.1% of VZs were entirely comprised of NPCs expressing the same color (**Figure 6D and 6E**), with the remainder consisting of NPCs with multiple distinct fluorescent barcodes (**Figure S6D**). To verify this was driven by clonal expansion of NPCs we examined markers of cell proliferation *in situ*. Ki67 and mitotic chromatin bodies were observed in Pax6^+^ VZ NPCs, confirming their proliferative phenotype, but not in postmitotic TBR1^+^ neurons (**Figures 6F and 6G**). Thus, by tracking barcoded NPCs as they differentiated, the rainbow reporter revealed most cortical VZs are derived from a single clonally dominant NPC.

## Discussion

A new era of single cell analyses, such as single cell transcriptomics and single cell epigenomics assays, has highlighted the wide range of molecular profiles that were previously unappreciated in bulk population assays (Angermueller *et al*., 2016; Nestorowa *et al*., 2016; Cusanovich *et al*., 2018; Nakanishi *et al*., 2019). These findings have been of special relevance to stem cell biology and directed differentiation approaches as they discovered transcriptional and epigenetic heterogeneity that were inferred to affect cellular functions, despite homogeneous expression of cell-type specific markers within the populations (Angermueller *et al*., 2016; Kumar, Tan and Cahan, 2017; Friedman *et al*., 2018). Here, we present a novel way to directly assess the phenotypic heterogeneity of individual PSCs and their differentiated derivatives within a population by labeling their membranes with unique colors and tracking them longitudinally by live cell imaging. This reporter is stably maintained through PSC differentiations and can also be induced with temporal control to label PSCs and their progeny in their differentiated or undifferentiated states. An array of cell behaviors can be extracted from the reporter including cell cycle kinetics, clonal expansion, apoptosis, cell morphology, and migration. We further demonstrate the longitudinal single cell lineage tracing and phenotyping afforded by this reporter can be supplemented with *in situ* molecular analyses of cell type and subcellular structures.

Tracking NPCs over a month of cortical neuron differentiation led to the surprise finding that clonally dominant progenitors self-aggregate to form complex cortical structures. The finding that monolayer cultures generate 3D cortical structures suggests the differentiation protocol robustly recapitulates *in vivo* signaling where the planar neural plate gives rise to the neural tube and its derivatives through morphogenesis (Yamaguchi *et al*., 2011). Longitudinal live imaging enabled determination of the timing of key neurogenesis events, such as VZ formation and neurite outgrowth. The membrane labels provided readouts for cell morphology, enabling clear determination of radial alignment and neurite formation. Though the underlying mechanisms remain incompletely understood, this analysis suggests clonal dominance results in VZs derived from a single NPC. Interestingly, unlike hiPSCs (**Figures 4E and 4F**) and NPCs expanding in media containing FGF (**Figure S5D**), NPCs cultured in neuron differentiation media without FGF displayed limited migration and dispersal. Consistent with this notion, the role of FGF in cell migration and early patterning of the neural primordium is well documented (Guillemot and Zimmer, 2011). While this warrants further study, the lack of NPC dispersal and mixing in differentiation media may contribute to the clonality of the VZs. In contrast to VZ NPCs, neurites from diverse lineages, expressing distinct color barcodes, intermixed and projected to distant VZs, suggesting neural networks form between VZs. Live imaging also demonstrated loss of clones over time, which may be related to NPC pruning in early neurogenesis (Yamaguchi *et al*., 2011). Although the dominance of a subset of NPCs resulting in clonal VZs was initially surprising to us, these findings are in line with neurogenesis in chick spinal cord. Studies using very sparse labeling (Leber and Sanes, 1995), and more recently rainbow labeling (Loulier *et al*., 2014), have demonstrated that discrete segments of the chick spinal cord are derived from individual progenitor cells that clonally expand and align around the VZ. Thus, this work provides the first evidence that similar clonality exists in models of human cerebral tissue. Since dysregulated NPC expansion can result in microcephaly or megacephaly (Chenn and Walsh, 2002; Lancaster *et al*., 2013; Johnson *et al*., 2018), understanding cellular mechanisms of NPC expansion on a per cell basis holds great potential to improve our understanding of neurodevelopmental disorders.

Previous works have utilized fluorescent reporters in PSCs (Gantz *et al*., 2012; Megyola *et al*., 2013) where a specific lineage is labeled by a single color. While it is possible to use these models to assess cell phenotypes, previous studies assessing clonal expansion with a single marker used less than 1% labeling to prevent neighboring cells to labeled by chance (Leber and Sanes, 1995; Miyaoka *et al*., 2012). This contrasts with our ability to discern the membrane of individual cells in highly migratory, dense cell populations of PSCs where greater than 90% of cells were labeled (**Video 4**). The range of color diversity and the persistence of the barcodes enabled tracking of individual cells over a week of culture even after cells had exponentially expanded and mixed (**Figures 3 and S5**).

Novel observations of hiPSC cell cycle kinetics were made with the rainbow reporter. Tracking individual cells over time revealed that hiPSCs from common ancestors had synchronized timing of cell divisions, consistent with PSC embryogenesis *in vivo* (O’Farrell, Stumpff and Tin Su, 2004; Deneke *et al*., 2016). This finding was undiscernible in previous *in vitro* studies of bulk PSC populations that concluded cell cycle arrest via serum deprivation or chemical inhibition of cell cycle machinery was required to achieve synchronized PSC divisions (Zhang *et al*., 2005; Yiangou *et al*., 2019). Though common ancestor cell division synchrony persists over several divisions, we note that, considering all cells in our study were derived from a single cell clone when we first generated the line, it appears that slight drift over time eventually leads to loss of synchrony. While less understood in humans, the extent of synchrony in the first few PSC divisions is predictive of embryo viability and implantation of *in vitro* fertilized human embryos (Wong *et al*., 2010; Meseguer *et al*., 2011; Desai *et al*., 2014), and the eventual loss of hiPSC division synchrony is consistent with studies in model organisms that demonstrated this synchronization persists for the first 13 rounds of cell division that give rise to the 8,192 cell embryo (O’Farrell, Stumpff and Tin Su, 2004). Future studies aimed at promoting PSC synchronization or isolating synchronized PSCs will help optimize differentiation and cell therapy strategies as cell division synchrony plays important roles in PSC cell type specification (Sela *et al*., 2012; Pauklin and Vallier, 2013; Yiangou *et al*., 2019) and improves the success of stem cell transplantation (Zhang *et al*., 2005; Desai *et al*., 2014).

We found this reporter of single cell lineage and clonal expansion had broader utility and enabled longitudinal quantification of cell morphology and migration. Studies have recently identified a subset population of PSCs that reside on the perimeter of PSC colonies (Li *et al*., 2010; Warmflash *et al*., 2014; Rosowski *et al*., 2015; Yoney *et al*., 2018; Nakanishi *et al*., 2019). These edge PSCs share hallmark properties with primitive endoderm and display distinct non-canonical Wnt signaling and gene expression that enhance their colony initiation capacity compared to PSCs localized in the colony interior (Yoney *et al*., 2018; Nakanishi *et al*., 2019). Their study found that PSCs isolated from the colony interior could acquire the edge-PSC signature 3 days after single cell dissociation and replating, however, it was unknown whether the edge subpopulation was stably localized to the colony perimeter. Our longitudinal study in intact PSC colonies found only a subset of PSCs persistently localize to the colony edge, while most PSCs at the edge rapidly migrate interiorly, suggesting the edge population is more dynamic and heterogeneous than previously appreciated. Interestingly, once PSCs exit the colony perimeter they very rarely return. This dynamic, one-way eviction from the colony periphery may be a mechanism that maintains colony shape as growing PSC colonies decrease the ratio of perimeter to internal cells (Peerani *et al*., 2007), which impacts PSC differentiation capacity (Peerani *et al*., 2007; Warmflash *et al*., 2014; Rosowski *et al*., 2015). In addition to labeling single cells, the reporter barcoded stem cell colonies, allowing unequivocal quantification of colony migration and tracking of colony fusion events. Morphological analysis showed hiPSCs become taller and skinnier as the culture became confluent. Interestingly, after the first day of culture we saw reductions in cell area that were not accompanied with increases in cell height, which we attribute to the presence of ROCK inhibitor in the first 26 hours of culture (Day0 and Day1 timepoints) which has been shown to promote cell spreading (Flevaris *et al*., 2007; Maldonado *et al*., 2016) without changing cell height (Nilius *et al*., 1999). Understanding the variability of these cellular phenotypes will be critical to standardize and efficiently manufacture stem cells for cell therapies (Trindade *et al*., 2017; Jossen *et al*., 2018).

The benefits of a multicolor reporter are accompanied with some limitations and special requirements. For complete avoidance of spectral bleed-through, specific microscope instrumentation (white light laser and spectral detector) (**Figure 1**) or fixation and staining of the samples (**Figure S2**) was needed, though native fluorescent protein signals could be distinguished with minimal spectral overlap using standard filters on our live imaging system (i.e. **Figure 3**). We note that not all hues were present in equal proportion, with the primary colors being most abundant, though unsurprisingly the chance of having more than one recombination event (i.e. mixed, non-primary colors) increased with higher Cre dose (**Figure 1**). Consistently, we saw most cells had at least one non-recombined rainbow cassette even in with high-dose (120 µg/mL) Cre based on staining for nFP (**Figure S2**), suggesting that color diversity will increase with higher Cre doses, although identification of dose that yields the greatest amount of color diversity without Cre toxicity is key for future studies (Jones *et al*., 2005; Cardozo *et al*., 2007; Schmidt-Supprian and Rajewsky, 2007). In addition, the nature of using multiple distinct fluorescent proteins presents challenges as they do not possess identical characteristics, such as protein folding time and extinction coefficients. We note that mKate2 expression was visible 12 hours after adding recombinant Cre, while the others were not present until 18 hours (*data not shown*). This is consistent with reports showing faster fluorophore maturation in mKate2 (Shcherbo *et al*., 2009). Despite efforts to reduce photobleaching, with a highly refined imaging protocol that showed no signs of phototoxicity, we still observe that the photostability of mOrange2 was slightly lower, which caused very subtle changes in color over the course of our time lapse imaging (**Video 2**, i.e. yellow eGFP^+^/mOrange2^+^ cells become slightly greener by the end of the time lapse). This is consistent with reported extinction coefficient values for the fluorescent proteins used (Cranfill *et al*., 2016).

Altogether, this rainbow PSC cell line has broad utility for studying cell phenotype over time and will enhance the field’s understanding of cell state in self-renewing PSC cultures and in directed differentiations to specific cell types. The integration of single cell molecular analysis with phenotypes revealed by this reporter opens new opportunities for forward genetics approaches that match observed proliferative and morphological phenotypes with specific molecular profiles. In conclusion, the rainbow PSC reporter line allows longitudinal tracking of individual cells and can uncover phenotypic heterogeneity that may improve human models of development, disease, drug discovery, and cell therapies.

## Experimental Procedures

### Contact for Reagents and Resource Sharing

Further information and requests for resources and reagents should be directed to and will be fulfilled by the Lead Contact, Jennifer Davis.

### PSC culture and differentiation

mTeSR Plus (STEMCELL technologies) was used for PSC maintenance culture (media changed daily) and was supplemented with 10µm Y27632 after passaging with StemPro Accutase (Gibco) onto Matrigel coated vessels. Recombinant CreTAT (Cre fused to cell permeability peptide, Excellgen eg1001) was added to media for 6 hours and the media was then changed. Previously established differentiation protocols were used, as detailed in the supplement. All experiments were performed with Passage 48-59 cells.

### Imaging and analysis

Images of native rainbow reporter fluorescence for spectral analysis were collect on a Leica SP8 confocal microscope with white light laser and spectral detector settings detailed in the supplement. Live imaging was performed on a Nikon TiE equipped with Yokogawa W1 spinning disk confocal system and high sensitivity EMCCD camera with configurations described in the supplement. Cells were manually traced and counted in ImageJ and cell area and centroid positions were determined using ImageJ’s “measure” function. Traced cells were manually tracked in time lapse experiments (as in **Video 4**) and the frame in which cytokinesis occurred for a given cell was recorded, with the time between frames being 12 minutes. Cell height was manually measured by analyzing z stacks taken with 0.4µm steps and the distance from the bottom to the top of the cell was calculated.

### Quantification and Statistical Analysis

GraphPad Prism V7 was used for One-Way ANOVAs, Two-Way ANOVAs, and Tukey posthoc tests. Standard One-way ANOVAs were performed for data comprising Figures 1, 2, 4A, 4B, 4D, S1, S3, S4C, and Repeat Measures ANOVA was used for Figure 3. Two-Way ANOVAs were performed for Figures 4H and S4B. P-value of <0.05 was considered significant. All average values described in the text and figures are mean values and error bars represent standard error of the mean. Each quantification was derived from a minimum of three independent (different passages) experiments.

## Acknowledgments

We would like to acknowledge Julie Mathieu and the University of Washington’s Institute for Stem Cell and Regenerative Medicine stem cell core, and Dale Hailey and Nathaniel Peters from the Garvey and Keck imaging cores. This work was supported by the following funding sources: NIH HL141187 & HL142624 (JD), NSF CMMI-1661730 (NJS), NIHDP2HL137188 (KRS), NIH F32HL143851 (DE), Washington Research Foundation postdoctoral fellowship (MCR), Gree Family Gift (JD/NJS/KRS/CEM), The University of Washington Ellison Foundation (JEY). CEM was supported by NIH grants R01HL128362, U54DK107979, R01HL128368, R01HL141570, R01HL146868, and a grant from the Fondation Leducq Transatlantic Network of Excellence.

## Author Contributions

Conceptualization, DE, CEM, NJS, JD; Methodology, DE, CEM, KRS, JEY, NJS, JD; Software, KMB; Investigation and Validation, DE, KS, KMB, RM, MCR, GWE; Formal Analysis, DE, KS, KMB, GWE; Writing – Original Draft; DE, NJS, JD; Writing – Review & Editing, DE, CEM, KRS, NJS, JD; Visualization, DE, KMB; Supervision, DE, CEM, NJS, JD; Funding Acquisition, DE, CEM, KRS, JEY, NJS, JD.

## References

Andäng, M. et al. (2008) ‘Histone H2AX-dependent GABA(A) receptor regulation of stem cell proliferation.’, Nature, 451(7177), pp. 460–4. doi: 10.1038/nature06488.

Angermueller, C. et al. (2016) ‘Parallel single-cell sequencing links transcriptional and epigenetic heterogeneity’, Nature Methods. 13(3), pp. 229–232. doi: 10.1038/nmeth.3728.

Bedogni, F. et al. (2010) ‘Tbr1 regulates regional and laminar identity of postmitotic neurons in developing neocortex’, PNAS. 107(29), pp. 13129–13134. doi: 10.1073/pnas.1002285107.

van Berlo, J. H. and Molkentin, J. D. (2014) ‘An emerging consensus on cardiac regeneration’, Nature Medicine. 20(12), pp. 1386–1393. doi: 10.1038/nm.3764.

Cai, D. et al. (2013) ‘Improved tools for the Brainbow toolbox.’, Nature methods, 10(6), pp. 540–7.

Cardozo, A. K. et al. (2007) ‘Cell-permeable peptides induce dose- and length-dependent cytotoxic effects’, Biochimica et Biophysica Acta (BBA) - Biomembranes. 1768(9), pp. 2222–2234. doi: 10.1016/J.BBAMEM.2007.06.003.

Chenn, A. and Walsh, C. A. (2002) ‘Regulation of cerebral cortical size by control of cell cycle exit in neural precursors’, Science, 297(5580), pp. 365–369. doi: 10.1126/science.1074192.

Cranfill, P. J. et al. (2016) ‘Quantitative assessment of fluorescent proteins.’, Nature methods. 13(7), pp. 557–62. doi: 10.1038/nmeth.3891.

Cusanovich, D. A. et al. (2018) ‘A Single-Cell Atlas of In Vivo Mammalian Chromatin Accessibility.’, Cell. 174(5), p. 1309–1324.e18. doi: 10.1016/j.cell.2018.06.052.

Deneke, V. E. et al. (2016) ‘Waves of Cdk1 Activity in S Phase Synchronize the Cell Cycle in Drosophila Embryos’, Developmental Cell. 38(4), pp. 399–412. doi: 10.1016/J.DEVCEL.2016.07.023.

Desai, N. et al. (2014) ‘Analysis of embryo morphokinetics, multinucleation and cleavage anomalies using continuous time-lapse monitoring in blastocyst transfer cycles.’, Reproductive biology and endocrinology : RB&E. 12, p. 54. doi: 10.1186/1477-7827-12-54.

Englund, C. et al. (2005) ‘Pax6, Tbr2, and Tbr1 are expressed sequentially by radial glia, intermediate progenitor cells, and postmitotic neurons in developing neocortex’, Journal of Neuroscience, 25(1), pp. 247–251. doi: 10.1523/JNEUROSCI.2899-04.2005.

Flevaris, P. et al. (2007) ‘A molecular switch that controls cell spreading and retraction.’, The Journal of cell biology, 179(3), pp. 553–65. doi: 10.1083/jcb.200703185.

Friedman, C. E. et al. (2018) ‘Single-Cell Transcriptomic Analysis of Cardiac Differentiation from Human PSCs Reveals HOPX-Dependent Cardiomyocyte Maturation’, Cell Stem Cell, 23(4), p. 586–598.e8. doi: 10.1016/j.stem.2018.09.009.

Gantz, J. A. et al. (2012) ‘Targeted genomic integration of a selectable floxed dual fluorescence reporter in human embryonic stem cells.’, PloS one. 7(10), p. e46971. doi: 10.1371/journal.pone.0046971.

Girshovitz, P. and Shaked, N. T. (2012) ‘Generalized cell morphological parameters based on interferometric phase microscopy and their application to cell life cycle characterization’, Biomedical Optics Express. 3(8), p. 1757. doi: 10.1364/BOE.3.001757.

Graham, V. et al. (2003) ‘SOX2 functions to maintain neural progenitor identity’, Neuron. 39(5), pp. 749–765. doi: 10.1016/S0896-6273(03)00497-5.

Guillemot, F. and Zimmer, C. (2011) ‘From cradle to grave: The multiple roles of fibroblast growth factors in neural development’, Neuron, pp. 574–588. doi: 10.1016/j.neuron.2011.08.002.

Hannan, N. R. F. et al. (2013) ‘Production of hepatocyte-like cells from human pluripotent stem cells’, Nature Protocols, 8(2), pp. 430–437. doi: 10.1038/nprot.2012.153.

Inoue, H. et al. (2014) ‘iPS cells: a game changer for future medicine.’, The EMBO journal, 33(5), pp. 409–17. doi: 10.1002/embj.201387098.

Johnson, M. B. et al. (2018) ‘Aspm knockout ferret reveals an evolutionary mechanism governing cerebral cortical size letter’, Nature. 556(7701), pp. 370–375. doi: 10.1038/s41586-018-0035-0.

Jones, S. W. et al. (2005) ‘Characterisation of cell-penetrating peptide-mediated peptide delivery.’, British journal of pharmacology. 145(8), pp. 1093–102. doi: 10.1038/sj.bjp.0706279.

Jossen, V. et al. (2018) ‘Manufacturing human mesenchymal stem cells at clinical scale: process and regulatory challenges’, Applied Microbiology and Biotechnology. 102(9), p. 3981. doi: 10.1007/S00253-018-8912-X.

Kocsisova, Z. et al. (2018) ‘Cell Cycle Analysis in the C. elegans Germline with the Thymidine Analog EdU.’, Journal of visualized experiments (140). doi: 10.3791/58339.

Kumar, P., Tan, Y. and Cahan, P. (2017) ‘Understanding development and stem cells using single cell-based analyses of gene expression.’, Development 144(1), pp. 17–32. doi: 10.1242/dev.133058.

Kusuma, S. and Gerecht, S. (2013) ‘Fast and furious: the mass and motion of stem cells.’, Biophysical journal, 105(4), pp. 837–8. doi: 10.1016/j.bpj.2013.07.021.

Lancaster, M. A. et al. (2013) ‘Cerebral organoids model human brain development and microcephaly’, Nature. 501(7467), pp. 373–379. doi: 10.1038/nature12517.

Leber, S. M. and Sanes, J. R. (1995) ‘Migratory paths of neurons and glia in the embryonic chick spinal cord’, Journal of Neuroscience, 15(2), pp. 1236–1248. doi: 10.1523/jneurosci.15-02-01236.1995.

Li, D. et al. (2010) ‘Integrated biochemical and mechanical signals regulate multifaceted human embryonic stem cell functions.’, The Journal of cell biology. 191(3), pp. 631–44. doi: 10.1083/jcb.201006094.

Livet, J. et al. (2007) ‘Transgenic strategies for combinatorial expression of fluorescent proteins in the nervous system’, Nature, 450(7166), pp. 56–62. doi: 10.1038/nature06293.

Lloyd, D. R. et al. (2000) ‘Relationship between cell size, cell cycle and specific recombinant protein productivity.’, Cytotechnology, 34(1–2), pp. 59–70. doi: 10.1023/A:1008103730027.

Loulier, K. et al. (2014) ‘Multiplex Cell and Lineage Tracking with Combinatorial Labels’, Neuron, 81(3), pp. 505–520. doi: 10.1016/j.neuron.2013.12.016.

Lundy, S. D. et al. (2013) ‘Structural and Functional Maturation of Cardiomyocytes Derived from Human Pluripotent Stem Cells’, Stem Cells and Development, 22(14), pp. 1991–2002. doi: 10.1089/scd.2012.0490.

Maldonado, M. et al. (2016) ‘ROCK inhibitor primes human induced pluripotent stem cells to selectively differentiate towards mesendodermal lineage via epithelial-mesenchymal transition-like modulation.’, Stem cell research, 17(2), pp. 222–227. doi: 10.1016/j.scr.2016.07.009.

Mallanna, S. K. and Duncan, S. A. (2013) ‘Differentiation of Hepatocytes from Pluripotent Stem Cells’, Current Protocols in Stem Cell Biology. pp. 1G.4.1-1G.4.13. doi: 10.1002/9780470151808.sc01g04s26.

Mariani, J. et al. (2012) ‘Modeling human cortical development in vitro using induced pluripotent stem cells.’, PNAS. 109(31), pp. 12770–5. doi: 10.1073/pnas.1202944109.

Megyola, C. M. et al. (2013) ‘Dynamic migration and cell-cell interactions of early reprogramming revealed by high-resolution time-lapse imaging.’, Stem cells 31(5), pp. 895–905. doi: 10.1002/stem.1323.

Meseguer, M. et al. (2011) ‘The use of morphokinetics as a predictor of embryo implantation’, Human Reproduction. 26(10), pp. 2658–2671. doi: 10.1093/humrep/der256.

Miyaoka, Y. et al. (2012) ‘Hypertrophy and Unconventional Cell Division of Hepatocytes Underlie Liver Regeneration’, Current Biology. 22(13), pp. 1166–1175. doi: 10.1016/J.CUB.2012.05.016.

Moreno, N. and González, A. (2017) ‘Pattern of neurogenesis and identification of neuronal progenitor subtypes during pallial development in xenopus laevis’, Frontiers in Neuroanatomy. S.A., 11. doi: 10.3389/FNANA.2017.00024.

Nakanishi, M. et al. (2019) ‘Human Pluripotency Is Initiated and Preserved by a Unique Subset of Founder Cells’, Cell. 177(4), p. 910–924.e22. doi: 10.1016/J.CELL.2019.03.013.

Nestorowa, S. et al. (2016) ‘A single-cell resolution map of mouse hematopoietic stem and progenitor cell differentiation.’, Blood. 128(8), pp. e20–31. doi: 10.1182/blood-2016-05-716480.

Nilius, B. et al. (1999) ‘Role of Rho and Rho kinase in the activation of volume-regulated anion channels in bovine endothelial cells.’, The Journal of physiology. 516 (Pt 1), pp. 67–74. doi: 10.1111/J.1469-7793.1999.067AA.X.

O’Farrell, P. H., Stumpff, J. and Tin Su, T. (2004) ‘Embryonic Cleavage Cycles: How Is a Mouse Like a Fly?’, Current Biology. 14(1), pp. R35–R45. doi: 10.1016/J.CUB.2003.12.022.

Orford, K. W. and Scadden, D. T. (2008) ‘Deconstructing stem cell self-renewal: genetic insights into cell-cycle regulation.’, Nature reviews. Genetics, 9(2), pp. 115–28. doi: 10.1038/nrg2269.

Paşca, A. M. et al. (2019) ‘Human 3D cellular model of hypoxic brain injury of prematurity’, Nature Medicine. doi: 10.1038/s41591-019-0436-0.

Pauklin, S. and Vallier, L. (2013) ‘The Cell-Cycle State of Stem Cells Determines Cell Fate Propensity’, Cell. 155(1), pp. 135–147. doi: 10.1016/J.CELL.2013.08.031.

Peerani, R. et al. (2007) ‘Niche-mediated control of human embryonic stem cell self-renewal and differentiation.’, The EMBO journal. 26(22), pp. 4744–55. doi: 10.1038/sj.emboj.7601896.

Rosowski, K. A. et al. (2015) ‘Edges of human embryonic stem cell colonies display distinct mechanical properties and differentiation potential.’, Scientific reports. 5, pp. 14218. doi: 10.1038/srep14218.

Rowe, R. G. and Daley, G. Q. (2019) ‘Induced pluripotent stem cells in disease modelling and drug discovery.’, Nature reviews. Genetics, 20(7), pp. 377–388. doi: 10.1038/s41576-019-0100-z.

Schmidt-Supprian, M. and Rajewsky, K. (2007) ‘Vagaries of conditional gene targeting’, Nature Immunology, 8(7), pp. 665–668. doi: 10.1038/ni0707-665.

Sela, Y. et al. (2012) ‘Human Embryonic Stem Cells Exhibit Increased Propensity to Differentiate During the G1 Phase Prior to Phosphorylation of Retinoblastoma Protein’, Stem Cells. 30(6), pp. 1097–1108. doi: 10.1002/stem.1078.

Shcherbo, D. et al. (2009) ‘Far-red fluorescent tags for protein imaging in living tissues.’, The Biochemical journal. 418(3), pp. 567–74. doi: 10.1042/BJ20081949.

Snippert, H. J. et al. (2010) ‘Intestinal Crypt Homeostasis Results from Neutral Competition between Symmetrically Dividing Lgr5 Stem Cells’, Cell. 143(1), pp. 134–144. doi: 10.1016/J.CELL.2010.09.016.

Trindade, F. et al. (2017) ‘Towards the standardization of stem cell therapy studies for ischemic heart diseases: Bridging the gap between animal models and the clinical setting’, International Journal of Cardiology, 228, pp. 465–480. doi: 10.1016/j.ijcard.2016.11.236.

Warmflash, A. et al. (2014) ‘A method to recapitulate early embryonic spatial patterning in human embryonic stem cells.’, Nature methods. 11(8), pp. 847–54. doi: 10.1038/nmeth.3016.

Wong, C. C. et al. (2010) ‘Non-invasive imaging of human embryos before embryonic genome activation predicts development to the blastocyst stage’, Nature Biotechnology. 28(10), pp. 1115–1121. doi: 10.1038/nbt.1686.

Yamaguchi, Y. et al. (2011) ‘Live imaging of apoptosis in a novel transgenic mouse highlights its role in neural tube closure’, Journal of Cell Biology. 195(6), pp. 1047–1060. doi: 10.1083/jcb.201104057.

Yiangou, L. et al. (2019) ‘Method to Synchronize Cell Cycle of Human Pluripotent Stem Cells without Affecting Their Fundamental Characteristics’, Stem Cell Reports. 12(1), pp. 165–179. doi: 10.1016/J.STEMCR.2018.11.020.

Yoney, A. et al. (2018) ‘WNT signaling memory is required for ACTIVIN to function as a morphogen in human gastruloids.’, eLife. doi: 10.7554/eLife.38279.

Zangle, T. A. et al. (2013) ‘Quantification of biomass and cell motion in human pluripotent stem cell colonies.’, Biophysical journal, 105(3), pp. 593–601. doi: 10.1016/j.bpj.2013.06.041.

Zhang, E. et al. (2005) ‘Cell cycle synchronization of embryonic stem cells: Effect of serum deprivation on the differentiation of embryonic bodies in vitro’, Biochemical and Biophysical Research Communications. 333(4), pp. 1171–1177. doi: 10.1016/J.BBRC.2005.05.200.

