## Supplemental text for "A Rainbow Reporter Tracks Single Cells and Reveals Heterogeneous Cellular Dynamics among Pluripotent Stem Cells and their Differentiated Derivatives"

**Supplemental Files**

**Videos 1-3, related to Figures 2, 4A, 4E, and 4F. Videos of 24 hour time-lapses of rainbow hiPSC cultures.**

**Video 4, related to Figures 2, 4A, 4E, and 4F. Video of 24-hour time lapse with cell traces.**

**Video 5, related to Figures 4C and 4D. Z-stack of hiPSCs showing height of cells at 4 days post plating, 0.4micron z-steps.**

**Video 6, related to Figure 5C. Rainbow cardiac myocytes beat in monolayer culture.**

**Video 7, related to Figure 6C and 6D. Optical sections and 3D reconstruction of clonal cortical structures, with DCX and Map2 staining shown in grayscale.**

**Supplemental File 1, related to Figure 1D. MATLAB code for spectral analysis.**

**Supplemental File 2, related to Figure 4E-4J. MATLAB code for migration analysis.**

### Supplemental Figure Legends

**Figure S1, related to Figure 1. Authentication of rainbow hiPSC model.**

(A) Confirmation of genotype showing presence or absence of WT allele, targeted knock-in allele, and additional incorporations in control and various rainbow-targeted clonal lines. (B) Copy number analysis of the number of cassettes in rainbow-targeted clonal lines. (C) Expanded view of Figure 1C showing individual channels and nuclei. Bar=20 µm. (D) Confirmation of the Tra-1-60 expression in rainbow hiPSCs. Bar=20 µm. (E) Karyotype analysis of Rainbow hiPSC cell before and after high dose treatment of Cre showing normal male karyotype.

**Figure S2, related to Figure 1. Visualization of spectral diversity and identification of unrecombined rainbow cassettes using immunostaining and conventional epifluorescence microscopy.**

(A) Widefield images of rainbow hiPSCs that were treated with various doses of Cre, immunostained, and imaged using conventional excitation/emission filter cubes. Note the GFP antibody recognizes both the fluorescent-membrane-localized eGFP as well as the nuclear-localized-non-fluorescent GFP mutant (nFP). Bar=20 µm. (B) Examples of hiPSCs expressing varying amounts and combinations of FPs after being treated with 120 µg/mL Cre and immunostained, including triple positive cells that lack nuclear localized nFP, indicating all rainbow cassettes were recombined. Each tile represents an area of 17.55µm x 14.62µm.

**Figure S3, related to Figure 1. Authentication of rainbow hESC model.**

(A) Representative confocal images of rainbow RUES2 human embryonic stem cells (hESCs) that were treated with various doses of Cre and imaged using white light laser and spectral detector settings that prevented spectral bleedthrough of the fluorescent proteins. Bar=20 µm. (B) Confirmation of genotype showing presence or absence of WT allele, targeted knock-in allele, and additional incorporations in control and rainbow-targeted clonal lines. (C) Copy number analysis of the number of cassettes in rainbow-targeted clonal lines. (D) Karyotype analysis of rainbow hESCs showing normal female karyotype.

**Figure S4, related to Figure 2. Rainbow reporter does not affect hiPSC proliferation or apoptosis**

(A) Immunostaining of phosphorylated-serine-10 of histone H3 (pH3, red) and Hoechst stain (green) at various timepoints post plating in wildtype isogenic control and transgenic hiPSCs with and without 120µg/mL Cre treatment. Bar=40 µm. (B) Quantification of pH3+ nuclei from panel A. (C) Quantification of TUNEL flow cytometry of samples harvested 72 hours after initial plating**.** (D) Example of apoptotic cell from time-lapse live cell imaging, bar=10 µm.

**Figure S5, related to Figure 5. Differentiation of rainbow labeled hiPSCs.**

(A) Timecourses for hepatoblast, cortical neuron, and cardiac myocyte differentiations. (B) Repeat imaging of NPC differentiation (Days 1-9), bar=200 µm, white arrows point to a clonally expanding cell shown in insets below. (C) Confirmation of NPC generation by Sox2 immunostaining. (D) Time course from NPC to neural rosette formation with example of clonally expanding and dispersing clone in the insets. (E) Brightfield image of isogenic wildtype control and rainbow-labeled rosettes, bar=200 µm. (F) 3D reconstruction (48 µm depth) of rosette structure showing combined rainbow fluorescence and a single channel. (G) Flow cytometry analysis of Day 14 cardiac myocytes using antibody against cardiac Troponin T or isotype control. (H) Unlabeled rainbow hiPSCs were differentiated to CMs, recombinant Cre was added on day 14, and cells were fixed and immunostained on day 24, bar=40 µm.

**Figure S6, related to Figure 6. Self-assembled 3D cortical structures are derived from clonally dominant progenitors.** (A) Rainbow signal showing clonally dominant NPC, viewed at the culture substrate (Z-bottom). The boxed region is shown at higher magnification at a Z-slice midway through the Z-stack (Z-middle). Panels below show maximum intensity projections of the same region after staining for several markers. (B) Confirmation of NPC marker expression in VZs. (A-B) Each coordinate grid represents a 500 x 500 µm square and each dot has a 10 µm diameter. (C) Maximum intensity projections of the rainbow signal and NPC and terminally differentiated cortical neuron markers, bar=50 µm. (D) Example of mixed VZ derived from multiple NPCs, showing Z-slices at 20 µm intervals, with 0 µm being at the culture substrate, bar=50 µm.

**Supplemental Experimental Procedures**

Generation of human PSC lines: We redesigned the brainbow3.2 construct (Addgene 45179), replacing the neuron-specific Thy1 promoter with a constitutive CAG promoter that is insensitive to cell-type and differentiation status (Gantz *et al.*, 2012; Oceguera-Yanez *et al.*, 2016). This construct was flanked with AAVS1 homology arms (Addgene 80490) to facilitate targeted knock-in using Gateway cloning (BP clonase Thermo Fisher 11789020 and LR clonase Thermo Fisher 11791020). We co-electroporated WTC11 hiPSCs and RUES2 embryonic stem cells with the resulting CAG-brainbow3.2 (Rainbow reporter) with AAVS1 homology arms construct and a plasmid expressing Cas9 and a guide RNA targeting the AAVS1 locus (Addgene 80494) using Amaxa Human Stem Cell Nucleofector (Lonza).

Characterization of rainbow PSC lines and validation of transgene: Single cells were expanded and screened by PCR (genotype) and qPCR (copy number) and we isolated genomic DNA for PCRs spanning the construct to confirm by Sanger sequencing (Eurofins).

*Genotyping primers:* Knockin allele 5' end: F- TCGACTTCCCCTCTTCCGATG, R- GAGCCTAGGGCCGGGATTCTC (1.2kb); knockin allele 3' end: F- GATGCCCGGCGTCTACTATG, R- CTCAGGTTCTGGGAGAGGGTAG (1.5kb); WT allele: F- TCGACTTCCCCTCTTCCGATG, R- CTCAGGTTCTGGGAGAGGGTAG (1.4kb); other insertions: F- GATGCCCGGCGTCTACTATG, R- GACCCAATATCAGGAGACTAGG (1.8kb) (other/additional insertions primers detect plasmid backbone and thus random insertion with the integration into the genome not mediated by homologous recombination where sequence outside of the homology arms get incorporated) (Ikeda *et al.*, 2018).

*Copy # qPCR primers:* Rainbow copy #: F- TTGACCTCGGAGGTGTAGTG, R- GACGGCGAGTTCATCTACAAG (191bp); internal control (HOXA7 genomic locus): F- CCGGCTCTTACGGCTACAAT, R- CCAAAGTGGCTGGAGGAGG (76bp).

*Sequencing primers* (we generated PCR products from genomic DNA template using primer pairs that spanned the rainbow construct and subsequently sequenced with the same primers): Primer pair 1: F- TCGACCGCCTTGATTCTCATGG, R- AGGACGACGGCAACTACAAGAC; primer pair 2: F- AGCAGGACCATGTGATCG, R- GACTACACCATCGTGGAACAG; primer pair 3: F- TGGGTCTAGATGCATTCG, R-GCCTCATCTACAACGTCAAG; primer pair 4: F- TGCTAGGGAGGTCGCAGTATC, R- CACATGGTCCTGCTGGAGTTC.

The number of unique hues was determined by copy number analysis demonstrating a total of four copies of the Brainbow cassette. With three possible fluorescent proteins that can be expressed from each cassette, the number of unique possible outcomes can be described by the trinomial theorem, $[(n+2)(n+1)]/2$, where $n$ = the number of copies of the cassette. To account for the possibility that a cell can undergo 1, 2, 3, or 4 recombination events this equation was solved for n=1, n=2, n=3, and n=4, and summed, indicating 34 unique recombination outcomes. However, twelve of the unique combinations result in duplicated colors that vary only by intensity (i.e. one copy of red versus four copies of red result in the same hue, though with a different signal intensity) and another four unique outcomes result in duplicated hues that vary only by tint (i.e. one copy of red versus one copy of green and one copy of blue with two copies of red results in the same red hue, though with a different tint of white). Thus, there are eighteen unique hues that can be represented on a color wheel. The primary colors were most represented, presumably due to the likelihood of only one Brainbow copy undergoing Cre-lox recombination and the greater proportion of recombination outcomes resulting in primary color expression.

Cell line karyotyping: Karyotyping analysis was performed on hiPSC and hESC lines by certified a cytologist Diagnostic Cytogenetics, Inc. (Seattle, WA) that indicated normal karyotypes.

Imaging: Images collected on a Leica SP8 confocal microscope used the following white light laser and spectral detector settings (excitation laser nm, emission spectral detector range nm-nm): AF 405 secondary/Hoechst (405, 410-500), eGFP (498, 504-540), mOrange2 (543, 558-584), mKate2 (590, 597-640). Antibody stains using conventional fluorescent secondary antibodies (AF405, AF 498, AF 555, AF 647) were imaged on a Nikon TiE widefield with standard DAPI, FITC, TRITC, and Cy5 filter cubes. Live imaging (24-hour time-lapses and long-term daily-imaging experiments) on a Nikon TiE equipped with Yokogawa W1 spinning disk confocal system and high sensitivity EMCCD camera was performed with the following settings (excitation fixed-LED laser nm, emission filter): eGFP (488, GFP), mOrange2 (561, TRITC), mKate2 (561, Cy5). For cell cycle kinetics 121 images were collected with 12 minutes between frames for each 24-hour experiment. For long-term daily live imaging experiments, we used culture slides that had gridded coordinate systems (Ibidi 80826-G500) to ensure re-imaging of the same cells/progeny of same ancestors, or 24 well plates that were physically marked with reference points (Ibidi 82406).

Spectral analysis: Images collected on a Leica SP8 confocal microscope with the described white light laser and spectral detector were analyzed on a per pixel basis and binned by the eighteen possible hues determine by genotype analysis using a custom generated MATLAB code (Supplemental File 1).

Migration analysis: Individual cells or cell colonies were manually traced, and the centroid positions relative to a fixed reference point were determine using ImageJ's "measure" function. The absolute positions or relative positions (where time 0 was set to X,Y coordinate 0,0) and migration were analyzed over time using custom MATLAB codes (Supplemental File 2). To quantify distances to the nearest colony edge, the distances from the most proximal cell membrane to the nearest colony edge was manually traced.

Staining: Samples were fixed with 4% PFA, permeabilized/blocked in 5% goat serum in PBS supplement with 0.1% TritonX-100, and stained with antibodies against: Oct4 (Abcam ab19857), Tra1 (Millipore MAB4360), cTnT (ThermoFisher MA5-12960), α-actinin (Sigma A7811), GFP (Kerafast EMU101), mCherry (recognizes mOrange2, Kerafast EMU105), TagRFP (recognizes mKate2, EMU107) phosphorylated-H3 (pH3, Abcam ab5176), Ki67 (BD Pharmigen 558615), Sox2 (Cell Signaling D6D9), Pax6 (DSHB), TBR1 (Abcam ab31940), DCX (Abcam ab18723), Map2 (Abcam ab92434), Ctip2 (Abcam ab18465), AFP (Agilent/Dako A000829), HNF4a (Abcam ab55223),. TUNEL (Roche 12156792910) and Hoechst (Life Technologies H3570) stains were performed according to manufacturers' guidelines.

Flow Cytometry: Cells were detached using Trypsin, followed by Trypsin inhibition with 10% fetal bovine serum (FBS), and in fixed with 4% PFA. The staining methods and reagents described above were used, modified for in suspension staining with cells centrifuged at 300g between each step. A Canto flow cytometer (BD Biosciences) was used and resulting data was analyzed following previously described procedures (Gerbin *et al.*, 2015).

Neural Differentiation: Neural differentiation of iPSCs into cortical excitatory neurons was carried out using established protocols with slight modifications (Mariani *et al.*, 2012; Rose *et al.*, 2018). A Basal Neural Maintenance Media [BNMM: 250mL DMEM/F12 + Glutamine (Gibco 11320-033), 250mL Neurobasal (Gibco 2110-049), 2.5mL N2 Supplement (Gibco 17502-048), 5mL B27 Supplement (Gibco 17504-044), 2.5mL GultaMax (Gibco 35050-061), 2.5mL ITS-A (Gibco 51300-044), 400uL BME (Thermo Fisher 21985023), and 2.5mL NEAA (Thermo Fisher 11140050)] was used throughout the protocol and supplemented with specific factors at the different stages of differentiation. iPSCs were dissociated with Accutase (Thermo Fisher Scientific # 00-455-56) for 4 minutes at 37ºC, washed 1 time with iPSC media and seeded onto Matrigel-coated 6-well plates at a density of 3M per well in iPSC media supplemented with 10uM Y-27632 ROCK Inhibitor (Ascent Scientific #ASC-129) overnight. The next day, neural induction was initiated using Neural Induction Media (NIM), by supplementing BNMM with 10uM SB431542 (Biogems #BG6675SKU301) and 0.5uM LDN 193189 (Biogems #BG5537SKU106). NIM was changed daily for 8 days. On day 8, cells were washed once with PBS, then treated with 1ml of warm Versene (Gibco #15040066) for 4 minutes at 37ºC. The Versene was aspirated off, then 3ml of NIM were added to the cells. A cell scraper was used to carefully scrape the cells off and a 5ml serological pipette was used to gently triturate the cells into small clumps. The cell clumps were transferred into new Matrigel-coated wells at a 1:3 split and incubated overnight in NIM plus 10µM Y-27632 ROCK inhibitor. The next day (day 9), the media was changed to BNMM and refreshed daily until well-defined neural rosettes were observed. At that point, Neural Stem Cell Media (NSCM) was fed to the cells and was made by supplementing BNMM with 10ug/ml FGF (R&D #233-FB/CF) and continued to day 16. On day 16, the cells were dissociated with Accutase, washed 1 time with BNMM and re-plated at a 1:3 split onto new Matrigel-coated wells in NSCM plus 10µM Y-27632 overnight. NSCM was refreshed daily up to day 23. On day 23, the cells were dissociated with Accutase, washed 1 time with BNMM and re-plated onto Matrigel-coated 10cm plates at a density of 4 million per 10cm plate or 125K cells per well of a 24-well plate in NSCM plus 10µM Y-27632 overnight. Neural differentiation into cortical neurons was initiated the next day by switching to ND+++ media, BNMM with 10ug/ml BDNF (PeproTech # 450-10), 10ug/ml GDNF (PeproTech # 450-02) and 250ug/ml dbcAMP (Sigma #D0627-1G). Half media changes were done every other day up to day 54, or longer, at which point the cells were fixed for ICC.

Cardiac Myocyte Differentiation: We used a previously established small molecule directed differentiation protocol with slight modification (Burridge *et al.*, 2014). Briefly, hiPSCs were seeded on Matrigel coated vessels (350,000 per well of 6-well plate or 100,000 per well of 24-well plate) in mTeSR media supplemented with 10µM Y-27632. 24 hours later media was changed to mTeSR media supplemented with 1µM Chiron 99021 (Cayman Chemicals). After 24 hours media was changed to RPMI (Gibco) supplemented with B27 supplement minus insulin (Thermo Fisher) and 5µM Chiron 99021. After 48 hours media was changed to RPMI supplemented with B27 supplement minus insulin and 2µM Wnt inhibitor Wnt-C59 (Selleckchem). After 48 hours media was changed to RPMI supplemented with B27 supplement plus insulin (cardios media), with recurring media changes every 2-3 days until endpoint. Single cell suspensions for analysis, replating and cardiac injections were generated by incubating the culture with a 1:1 mixture of 0.25% Trypsin and Versene for 7 minutes at 37C. Digestion was stopped with Stop Solution (cardios media + 10% FBS + 0.2 Kunitz units/µL DnaseI) and cells were replated in cardios media supplemented with 10µM Y-27632 and 5% FBS for the first 24 hours, then returned to standard Cardios media and replaced every 2-3 days.

Hepatic Differentiation: Rainbow labeled hiPSCs were differentiated to hepatic progenitors using previously published methods (Mallanna and Duncan, 2013). Briefly, hiPSCs were seeded at 400,000 cells/well in 24-well plate format in mTeSR1 containing 10µM Y-27632 and allowed to adhere overnight. The following morning (17 hr later) media was changed to mTeSR1 without Y-27632. Following an additional 8hr of culture in mTeSR1, media was changed to RPMI-1640 (RPMI, Gibco) containing 1X B-27 Supplement minus insulin (ThermoFisher), 0.5X non-essential amino acids (NEAA, ThermoFisher), 1X Penicillin/Streptomycin, 100ng/ml Activin A (R&D Systems), 10ng/ml BMP4 (R&D Systems), and 20ng/ml bFGF (Invitrogen) for two days with daily media changes (Differentiation days one and two). On day three media was changed to RPMI containing 1X B-27 Supplement minus insulin, 0.5X NEAA, 1X Penicillin/Streptomycin, and 100ng/ml Activin A for three days with daily media changes. Cells were cultured in a 5% $CO_2$, 37°C incubator with ambient $O_2$ for days 0-5. On day six media was changed to RPMI containing 1X B-27 Supplement (containing insulin, ThermoFisher), 0.5X NEAA, 1X Penicillin/Streptomycin, 20ng/ml BMP4, and 10ng/ml bFGF for five days with daily media changes and cultures were moved from ambient $O_2$ to 4% $O_2$, 5% $CO_2$, 37°C for the remainder of the differentiation. On day 11 media was changed to RPMI containing 1X B-27 Supplement, 0.5X NEAA, 1X Penicillin/Streptomycin, and 20ng/ml HGF (Peprotech) for two days with daily media changes.

**Supplemental References**

Burridge, P. W. *et al.* (2014) ‘Chemically defined generation of human cardiomyocytes’, *Nature Methods*. Nature Publishing Group, 11(8), pp. 855–860. doi: 10.1038/nmeth.2999.

Gantz, J. A. *et al.* (2012) ‘Targeted genomic integration of a selectable floxed dual fluorescence reporter in human embryonic stem cells.’, *PloS one*. Edited by M. Daadi, 7(10), p. e46971. doi: 10.1371/journal.pone.0046971.

Gerbin, K. A. *et al.* (2015) ‘Enhanced Electrical Integration of Engineered Human Myocardium via Intramyocardial versus Epicardial Delivery in Infarcted Rat Hearts’, *PLoS ONE*. Public Library of Science, 10(7), p. e0131446. doi: 10.1371/JOURNAL.PONE.0131446.

Ikeda, K. *et al.* (2018) ‘Efficient scarless genome editing in human pluripotent stem cells’, *Nature Methods*. Nature Publishing Group, 15(12), pp. 1045–1047. doi: 10.1038/s41592-018-0212-y.

Mallanna, S. K. and Duncan, S. A. (2013) ‘Differentiation of Hepatocytes from Pluripotent Stem Cells’, in *Current Protocols in Stem Cell Biology*. Hoboken, NJ, USA: John Wiley & Sons, Inc., p. 1G.4.1-1G.4.13. doi: 10.1002/9780470151808.sc01g04s26.

Mariani, J. *et al.* (2012) ‘Modeling human cortical development in vitro using induced pluripotent stem cells.’, *Proceedings of the National Academy of Sciences of the United States of America*. National Academy of Sciences, 109(31), pp. 12770–5. doi: 10.1073/pnas.1202944109.

Oceguera-Yanez, F. *et al.* (2016) ‘Engineering the AAVS1 locus for consistent and scalable transgene expression in human iPSCs and their differentiated derivatives’, *Methods*, 101, pp. 43–55. doi: 10.1016/j.ymeth.2015.12.012.

Rose, S. E. *et al.* (2018) ‘Leptomeninges-Derived Induced Pluripotent Stem Cells and Directly Converted Neurons From Autopsy Cases With Varying Neuropathologic Backgrounds.’, *Journal of neuropathology and experimental neurology*. Oxford University Press, 77(5), pp. 353–360. doi: 10.1093/jnen/nly013.
