## Supplementary figures and images for "A Rainbow Reporter Tracks Single Cells and Reveals Heterogeneous Cellular Dynamics among Pluripotent Stem Cells and their Differentiated Derivatives"

### Supplemental figures

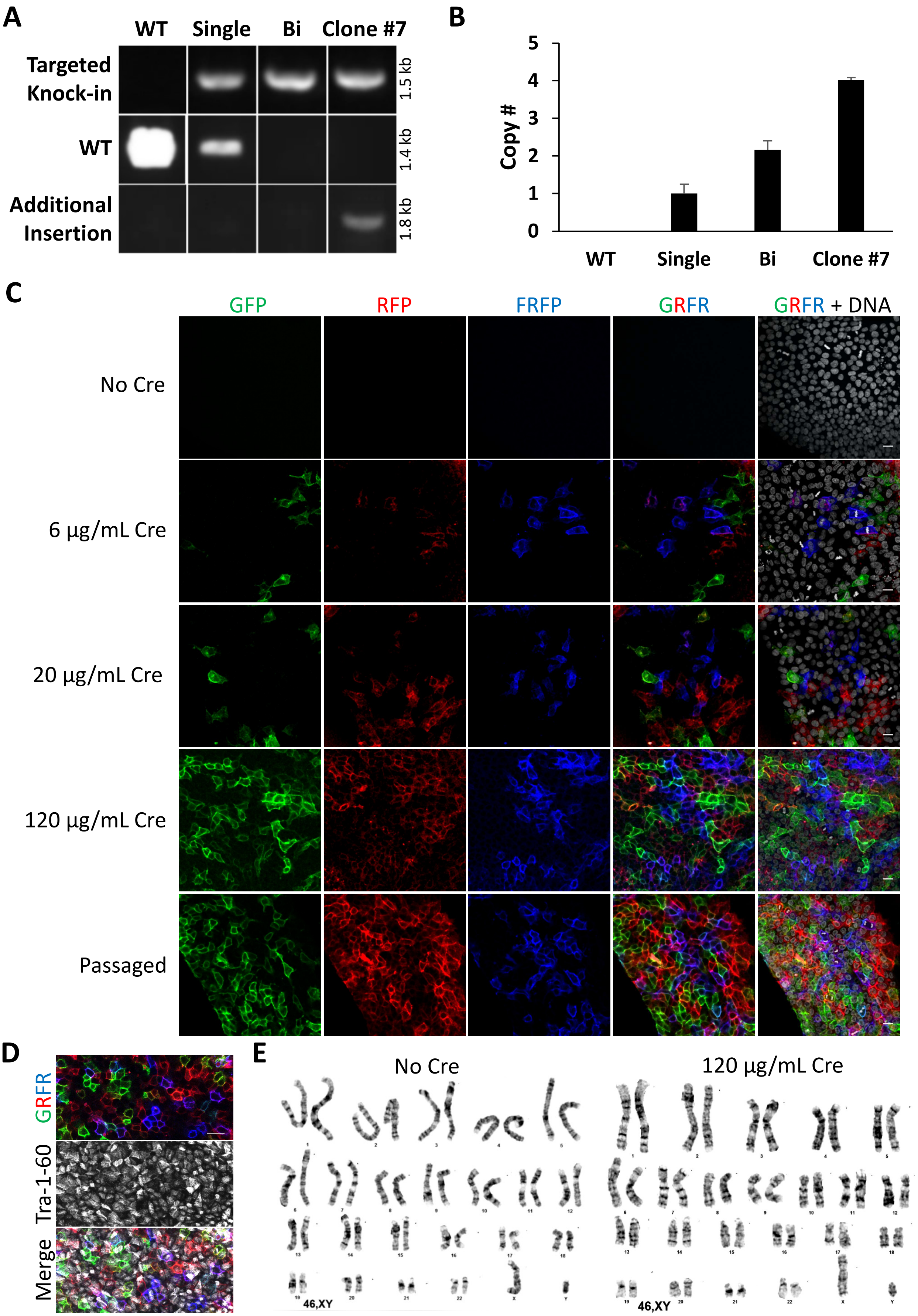

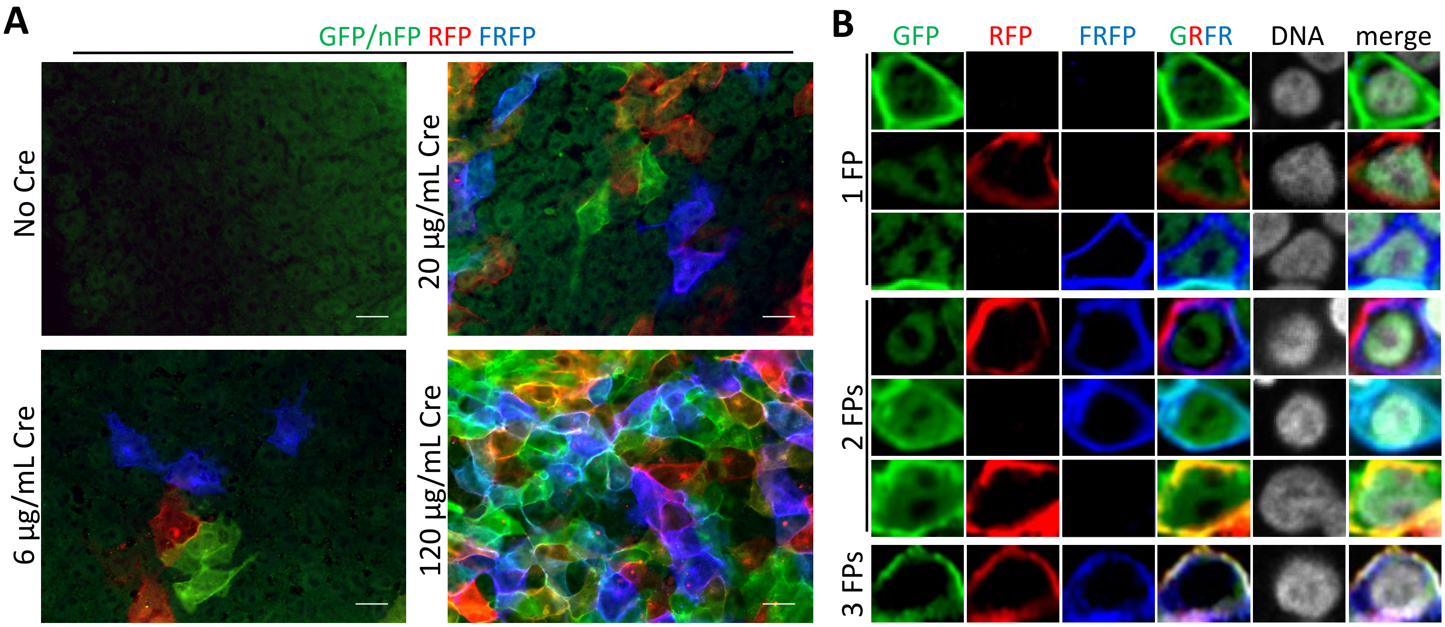

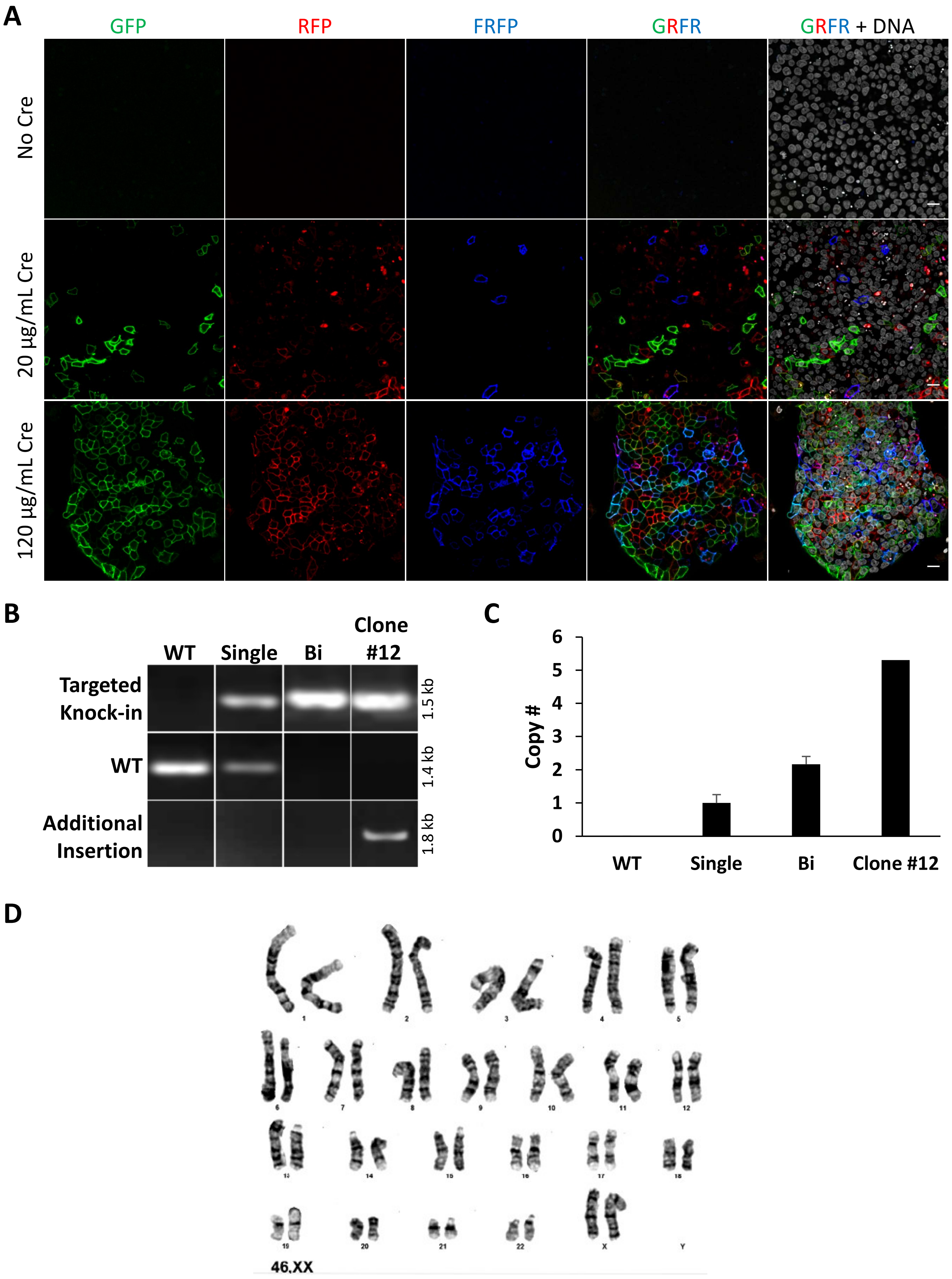

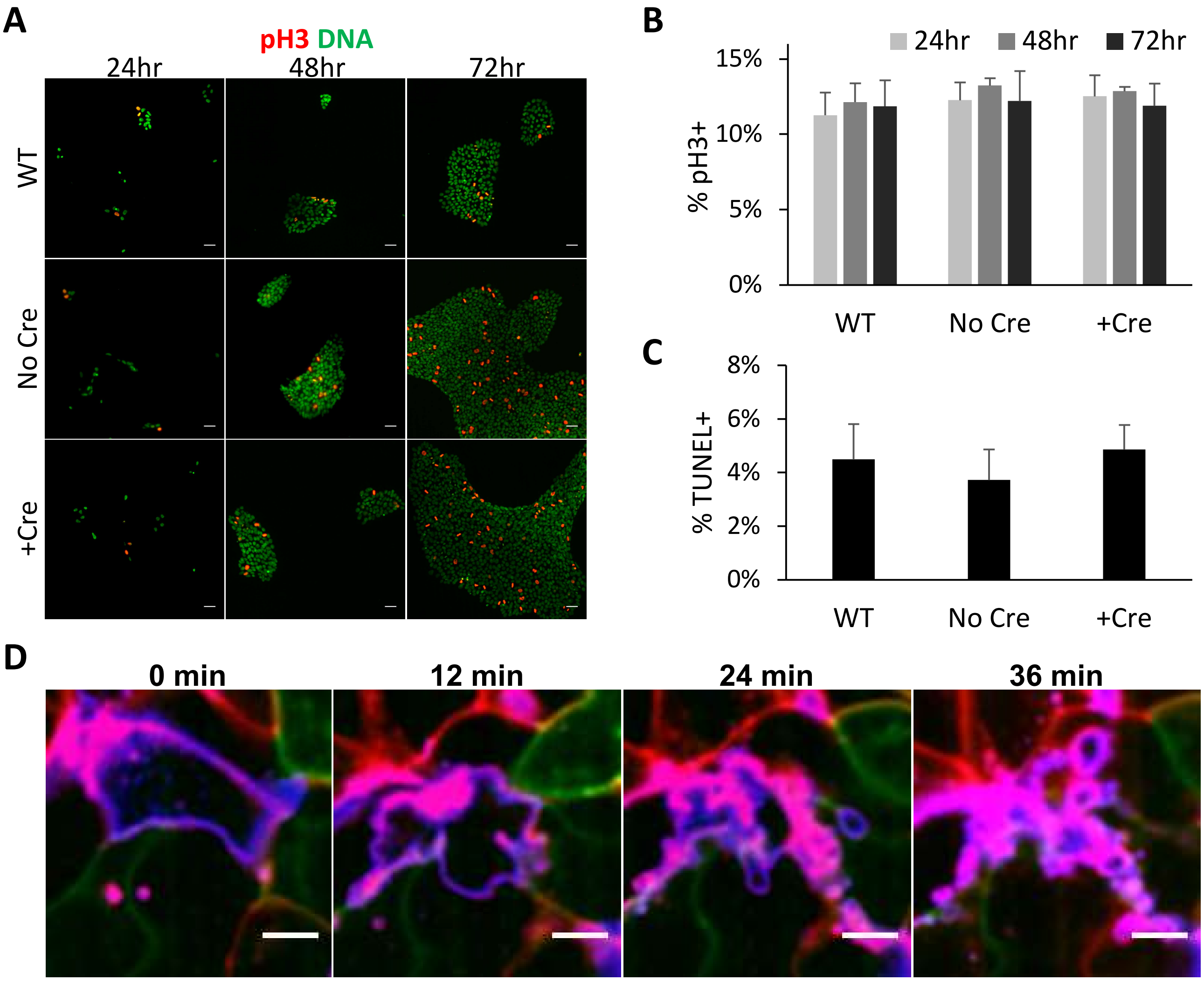

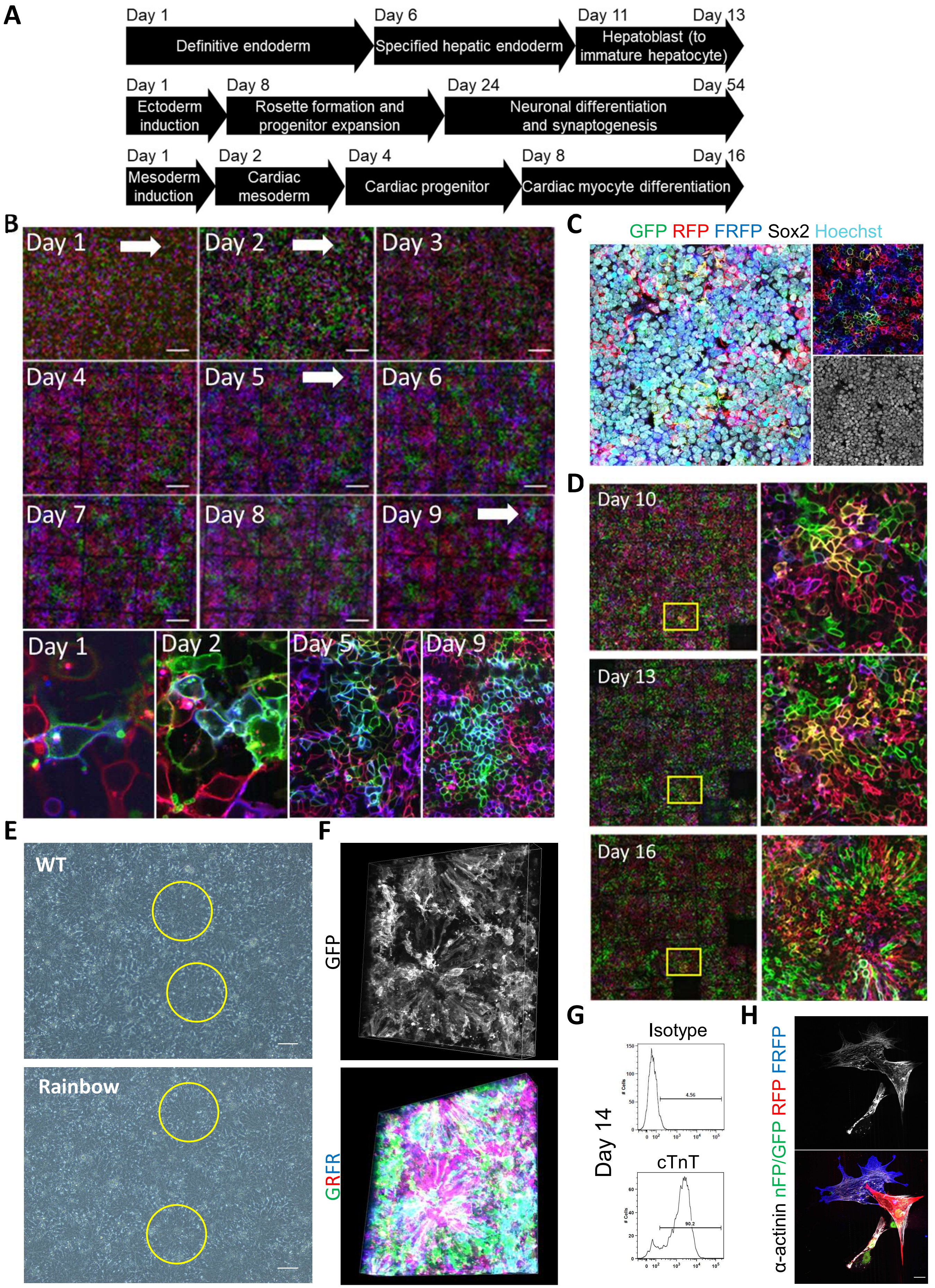

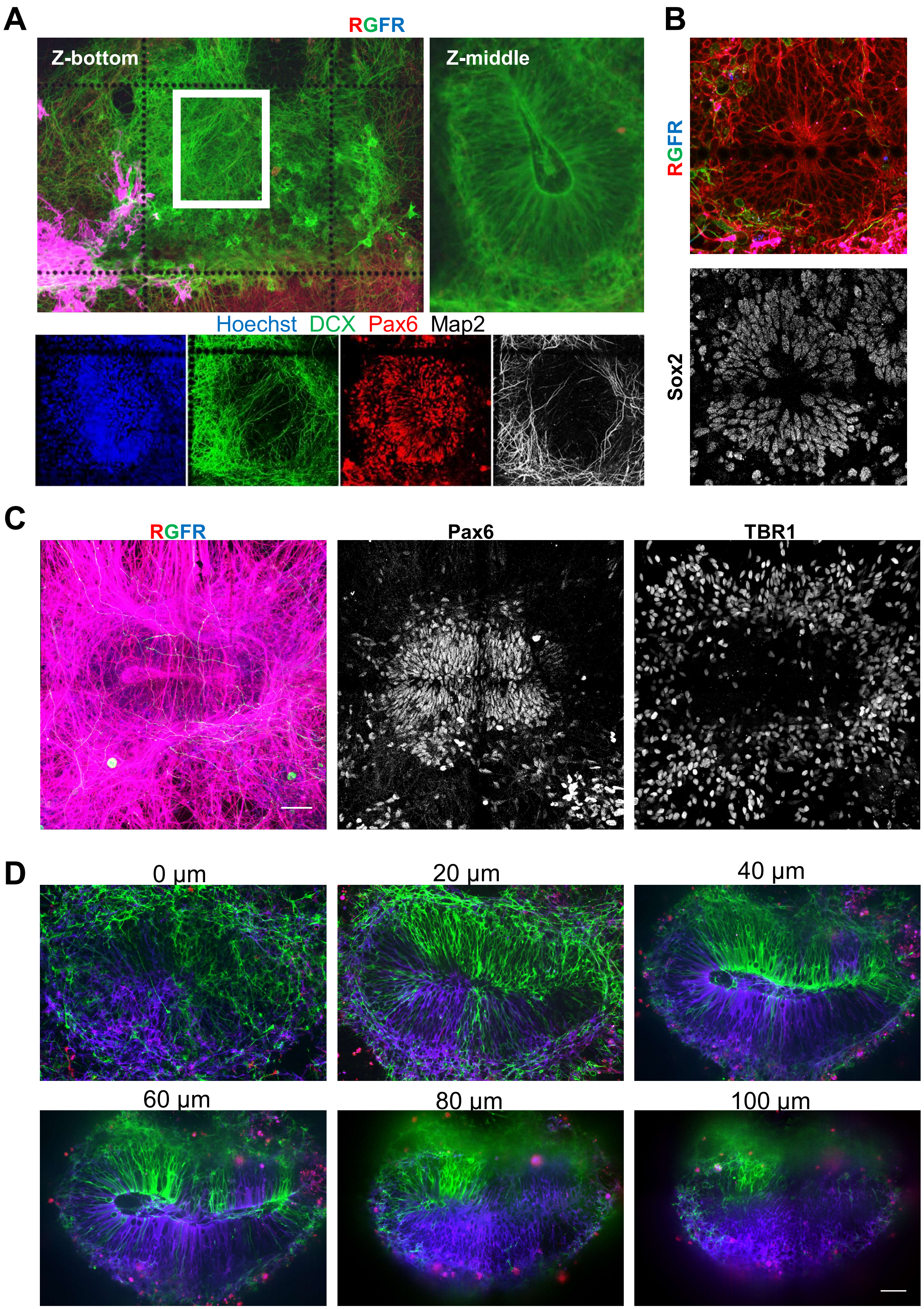
